# Genomic analyses of 10,376 individuals provides comprehensive map of genetic variations, structure and reference haplotypes for Chinese population

**DOI:** 10.1101/2021.02.06.430086

**Authors:** Peikuan Cong, Wei-Yang Bai, Jinchen Li, Nan Li, Sirui Gai, Saber Khederzadeh, Yuheng Liu, Mochang Qiu, Xiaowei Zhu, Pianpian Zhao, Jiangwei Xia, Shihui Yu, Weiwei Zhao, Junquan Liu, Penglin Guan, Yu Qian, Jianguo Tao, Mengyuan Yang, Geng Tian, Shuyang Xie, Keqi Liu, Beisha Tang, Hou-Feng Zheng

## Abstract

Here, we initiated the Westlake BioBank for Chinese (WBBC) pilot project with 4,535 whole-genome sequencing individuals and 5,481 high-density genotyping individuals. We identified 80.99 million SNPs and INDELs, of which 38.6% are novel. The genetic evidence of Chinese population structure supported the corresponding geographical boundaries of the Qinling-Huaihe Line and Nanling Mountains. The genetic architecture within North Han was more homogeneous than South Han, and the history of effective population size of Lingnan began to deviate from the other three regions from 6 thousand years ago. In addition, we identified a novel locus (*SNX29*) under selection pressure and confirmed several loci associated with alcohol metabolism and histocompatibility systems. We observed significant selection of genes on epidermal cell differentiation and skin development only in southern Chinese. Finally, we provided an online imputation server (https://wbbc.westlake.edu.cn/) which could result in higher imputation accuracy compared to the existing panels, especially for lower frequency variants.

## Introduction

Understanding the architecture of the human genome has been a fundamental approach to precision medicine. Over the past decade, great progress has been made to unravel either the genetic basis of complex traits and diseases (Timpson et al., 2018) or the human evolutionary history (Nielsen et al., 2017). The in-depth analysis of global populations with diverse ancestry could improve the understanding of the relationship between genomic variations and human diseases (2019). However, genetic studies exhibited a vast imbalance in global population, with individuals of European descent took up ∼79% of all genome wide association study (GWAS) participants (2019; Martin et al., 2019). Similarly, most of the whole-genome sequencing (WGS) efforts were predominantly conducted on European populations, such as Dutch (Genome of the Netherlands, 2014), UK (Consortium et al., 2015) and Icelandic population (Gudbjartsson et al., 2015). Even in larger genomic projects such as the Trans-Omics for Precision Medicine (TOPMed) program, which consisted of ∼155k participants from >80 different studies, only 9% of samples were of Asian descent (Kowalski et al., 2019). Therefore, large-scale genomic data are required to understand the genetic basis in Asian population. Recently, some studies have sequenced and analyzed the Asian populations including Japanese (Nagasaki et al., 2015) and Korean (Jeon et al., 2020). The Singapore SG10K pilot project reported 4,810 whole-genome sequenced samples, including 903 Malays, 1,127 Indians and 2,780 Chinese (Wu et al., 2019), and the pilot study of the GenomeAsia 100K Project presented a dataset of 1,267 individuals from different countries across Asia (GenomeAsia, 2019).

China, as the most populated country, is a multi-ethnic nation, in which the Han Chinese accounts for 90% of the population. Generally, the entire territory of the country (34 administrative divisions, including provinces, municipalities and special administrative regions) could be divided into Northern and Southern area by the geographical barrier of the Qinling-Huaihe Line (Shi et al., 2019). The Qinling Mountains are the east-west mountain range that stretch across the south of Gansu and Shaanxi provinces. The ∼1,000 kilometers long Huaihe River flows through the south of Henan province and the middle of Anhui and Jiangsu provinces. To some extent, the climate, culture, lifestyle and cuisine between the Northern and Southern regions were differed. Lingnan area is the region in the south of Nanling mountains (with five ridges) and the southeast of Yunnan-Guizhou Plateau in southern China, which refers to the administrative divisions of Guandong, Guangxi, Hainan, Hong Kong and Macao (Xie et al., 2020). Although the genetic structure of north-south differentiation in the Chinese population was consistently observed in previous studies (Cao et al., 2020; Chen et al., 2009; Chiang et al., 2018; Liu et al., 2018; Xu et al., 2009), clear subgrouping of the population was not always consistent. For example, Cao et al distinguished the Han Chinese into 7 population subgroups (Cao et al., 2020), while Xu et al clustered the Han Chinese into 3 sub-areas (Xu et al., 2009).

Despite the above efforts, the Chinese population was still underrepresented in human genetic studies, which could increase the health disparities if Chinese personal genomes were underserved (Martin et al., 2019; Popejoy and Fullerton, 2016; Sirugo et al., 2019). In addition, our previous study (Bai et al., 2019) demonstrated that, even with the Haplotype Reference Consortium (HRC) reference panel which contained 64,976 human haplotypes (McCarthy et al., 2016), the imputation of Chinese population could not reach the highest accuracy, a population specific reference panel was still needed (Bai et al., 2019). Therefore, the genetic study of Chinese population has the potential to benefit ∼20% of the world population, and provide a comparison to the rest of the world. Thus, we initiated the Westlake BioBank for Chinese (WBBC) project (Zhu et al., 2020) to characterize the genomic variation and population structure in a large-scale cohort aiming to collect ∼100,000 samples with deep phenotypes. Here, the findings of the pilot project of the WBBC from 10,376 samples were described, covering 29 out of 34 administrative divisions of China.

## Results

### The WBBC Pilot Dataset and Variants Identified

The WBBC pilot project sampled 10,376 individuals from 29 of 34 administrative divisions of the People’s Republic of China (Figure 1A and Table S1). We performed whole genome sequencing (WGS) in 4,535 individuals on NovaSeq 6000 platform. Hunan (n = 3,203) and Jiangxi (n = 719) provinces accounted for 87% of all WGS samples. After removing contaminated and duplicated samples, 4,489 unrelated individuals were retained for downstream analyses and statistics. The mean sequencing coverage was 13.9 ×, which covered 99.77% of the genome, with a range between 9.6 × and 65.2 × (Figure S1 and Table S2). Additionally, 5,841 individuals were genotyped by high-density Illumina Asian Screening Array (ASA) with 73.9M variants. Shandong (n = 2,801) and Jiangxi (n = 1,730) provinces comprised 77.6% of all ASA genotyped samples.

**Figure 1.**
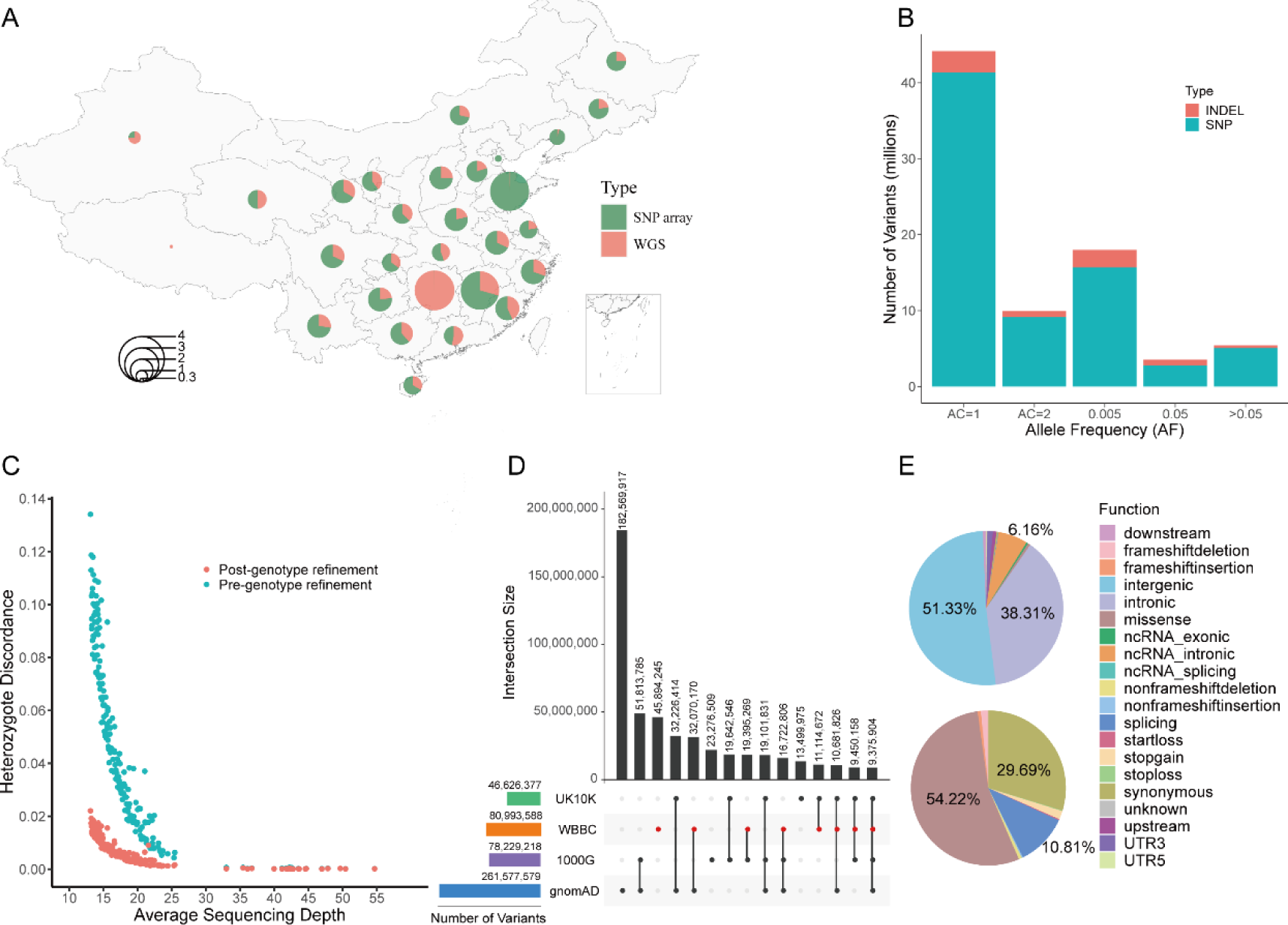
The statistics of samples and variants in the WBBC-cohort. (A) Sample distribution and statistics by geography. The pie chart shows the number of samples in each administrative division. The proportion of samples sequenced by whole-genome sequencing (WGS) and those genotyped by high-density Illumina Asian Screening Array (ASA) were marked in red and green, respectively. The values were converted into Log10. (B) The number of SNV and INDEL variants identified in the WBBC cohort in six frequency bins: AC = 1, AC = 2, AC > 2 & AF < 0.005, 0.005 ≤ AF ≤ 0.05, and AF > 0.05. (C) The estimated heterozygote discordance rate versus sequencing depth for 184 samples. The red dots indicate the average proportion of post-genotype refinement via Beagle tools and the green dots denote the raw genotype calls. (D) The number of variants in 22 autosomes and X chromosome in the WBBC, 1000 Genome Project (1000G), gnomAD, and UK10K datasets. The horizontal bar plot shows the total number of variants in each of the four datasets. The individual dots and connected dots indicate each dataset and a combination of two or more datasets, respectively. Each vertical bar represents the number of variants in each dataset or overlapping variants in those datasets. (E) Functional annotations of all novel variants were absent in dbSNP Build 151. The proportion of each category was filled with a different color. The upper pie chart showed all the 20 classification terms. The bottom pie chart only displayed the variants in the coding and splice regions (10bp from exon-intron boundary).

Using BWA (Li and Durbin, 2010) and GATK4 (Van der Auwera et al., 2013) recommendation pipeline and analysis strategy (Figure S2), we identified 80,993,588 variants after filtration from 104.2 million total raw variants (Ts/Tv = 2.15), including 73,942,614 single-nucleotide variants (SNVs) and 7,050,974 insertions and deletions (INDELs, < 50bp in length) (Figure S3A). Of these, 93.3% comprised rare (allele frequency, AF < 0.5%) and low-frequency (AF = 0.5-5%) variants, with the majority of variants being singletons (44.2 million, 54.5%, Figure 1B and Table S3). Additionally, C>T, G>A, A>G and T>C constituted the most common SNV substitution types (Figure S4), and the length of INDELs mainly distributed between −10bp and 10bp (Figure S5). We provided a database of genetic variations for the Han population in four sub-regions (North, Central, South and Lingnan) (https://wbbc.westlake.edu.cn/genotype.html).

To assess the SNV variants calling accuracy and sensitivity, 184 individuals from the whole genome sequencing samples (13.3 × - 54.7 ×) were also genotyped on ASA chip. After filtration, about 0.5 million common variants in autosomes detected by both WGS and SNP array were used to estimate the genotype concordance. We also performed LD-based genotype refinement via BEAGLE tools (Browning et al., 2018). Supposing the genotype from SNP arrays as the reference allele, the non-reference genotype concordance rate extended to 99.88% at 25 × with increasing sequencing depth (Figure S6A). The non-reference sensitivity and specificity had an effective increase after genotype refinement with BEAGLE from 0.9211 to 0.9924 and from 0.9931 to 0.9999, whereas it had inconspicuous improvement on homozygote genotype concordance (Figures S6B, S6C and S6D). The heterozygote discordance rate of genotypes was reduced 6-fold from 0.134 to 0.022 at 13.1 × sequencing depth and 4-fold from 0.004 to 0.001 at 25.3 × after genotype refinement (Figure 1C).

### Novel Variants and Functional Annotation

Comparing the variants with the WBBC and other existing databases, 45,894,245 variants were found not to present in the 1000 Genome Project (1KG) (Genomes Project et al., 2015), gnomAD (Lek et al., 2016) and UK10K (Consortium et al., 2015) (Figure 1D). Of these, 45.84 (99.878%) million were rare variants (MAF < 0.005), 45,093 (0.098%) were low frequency variants (0.005 ≤ MAF ≤ 0.05) and 10,836 (0.024%) were common variants (MAF > 0.05). We found 31.27 (38.6%) million novel variants that were not present in dbSNP Build 151 (Sherry et al., 2001), including 28,969,267 (92.6%) SNVs and 2,301,842 (7.4%) INDELs (Figure S3B). Of these variants, singletons accounted for 83.3%, and 99.95% of the variants (31.26 million) were rare with MAF < 0.005.

To characterize variants with a biological consequence, we annotated all the variants by ANNOVAR tools (Wang et al., 2010). As expected, 77,377,200 (95.5%) variants were in intergenic and intronic regions (Table S3). The variants in intergenic and intronic regions comprised 89.64% of novel variants (Figure 1E). In coding and splice regions, the missense accounted for 54.22% of the novel variants, while synonymous and splice variants made up 40.5% of the novel variants (Figure 1E). We also found that the missense, stop-gain, frameshift indels and non-frameshift indels variants were markedly increased among rare variants, compared with low frequency and common variants, which were signatures of population expansion and weak purifying selection (Figure S7). We predicted about 300,000 deleterious variants by SIFT (Kumar et al., 2009), PolyPhen-2 (Adzhubei et al., 2013) or MutationTaster (Schwarz et al., 2014) in 4,489 individuals, with the majority of variants being rare alleles with MAF < 0.5% (Table S3). Interestingly, we also identified 1,842 pathogenic or likely pathogenic variants recorded by ClinVar (Landrum et al., 2014) in our dataset. Of these predicted disease-causing variants, 97.4% variants were rare, 1.7% variants were low frequency, and 0.9% were common variants, which arose from selection pressure subjected on these rare variants.

### Variants in Individual Genome

We selected 1,151 healthy individuals for the autosomal variants’ statistic of a personal genome. On average, an individual carried 2,936,012 SNVs and 191,333 INDELs, including 8,915 missense, 10 stop loss, 70 stop gain and 126 frameshift or non-frameshit indels (Table 1). In total, 96.5% of the variants were located in intergenic and intronic regions. For the disease-associated variants predicted in silico, about 1,623 variants were deleterious by SIFT (Kumar et al., 2009), 1,714 variants were probably or possibly damaging by Polyphen2 (Adzhubei et al., 2013), and 8,591 variants were disease-causing by MutationTaster (Schwarz et al., 2014). In these pathogenic and deleterious variants, we observed the higher ratio of heterozygote and non-reference homozygote (Het/Hom) (Table 1). The proportions of Het/Hom were also very high in novel SNVs and INDELs variants, which indicated that the majority of novel variants occurred as heterozygotes in the Han Chinese population.

**Table 1.** The average number of autosomal variants in each genome of four regions.

|  | **Han Chinese**<br>(n = 1151,<br>Depth = 16.65×) |  | **North**<br>(n = 213,<br>Depth = 16.56×) |  | **Central**<br>(n = 42,<br>Depth = 16.67×) |  | **South**<br>(n = 845,<br>Depth = 16.66×) |  | **Lingnan**<br>(n = 51,<br>Depth = 16.89×) |  |
| --- | --- | --- | --- | --- | --- | --- | --- | --- | --- | --- |
| Annotation | No. | Het/Hom | No. | Het/Hom | No. | Het/Hom | No. | Het/Hom | No. | Het/Hom |
| **Type** |  |  |  |  |  |  |  |  |  |  |
| SNVs | 2,936,012 | 1.27 | 2,942,027 | 1.28 | 2,935,356 | 1.26 | 2,934,641 | 1.26 | 2,934,137 | 1.26 |
| Novel SNVs (not in dbSNP151) | 7,855 | 346.65 | 8,876 | 422.91 | 8,544 | 392.03 | 7,602 | 326.45 | 7,212 | 361.04 |
| Novel SNVs (not in 1000G) | 222,462 | 1.52 | 223,170 | 1.54 | 222,604 | 1.52 | 222,298 | 1.51 | 222,107 | 1.51 |
| INDELs | 191,333 | 2.87 | 191,636 | 2.86 | 191,214 | 2.84 | 191,212 | 2.87 | 192,163 | 2.87 |
| Novel INDELs (not in dbSNP151) | 1,453 | 82.6 | 1,497 | 85.05 | 1,486 | 85.07 | 1,440 | 81.79 | 1,452 | 83.74 |
| Novel INDELs (not in 1000G) | 79,853 | 5.93 | 79,342 | 5.86 | 79,593 | 5.87 | 79,953 | 5.95 | 80,556 | 5.96 |
| **Location** |  |  |  |  |  |  |  |  |  |  |
| Exon | 31,419 | 1.36 | 31,480 | 1.38 | 31,401 | 1.36 | 31,408 | 1.36 | 31,356 | 1.35 |
| Intron | 1,335,361 | 1.34 | 1,337,888 | 1.35 | 1,334,826 | 1.34 | 1,334,788 | 1.34 | 1,334,732 | 1.33 |
| Splice-site | 3,221 | 1.44 | 3,223 | 1.46 | 3,212 | 1.43 | 3,220 | 1.44 | 3,219 | 1.44 |
| UTR | 35,272 | 1.36 | 35,355 | 1.38 | 35,327 | 1.36 | 35,247 | 1.35 | 35,281 | 1.36 |
| Upstream | 19,367 | 1.3 | 19,412 | 1.32 | 19,382 | 1.31 | 19,356 | 1.30 | 19,353 | 1.29 |
| Downstream | 20,326 | 1.37 | 20,382 | 1.39 | 20,369 | 1.37 | 20,309 | 1.36 | 20,330 | 1.36 |
| Intergenic | 1,682,380 | 1.31 | 1,685,921 | 1.33 | 1,682,052 | 1.31 | 1,681,525 | 1.31 | 1,682,029 | 1.30 |
| **Function** |  |  |  |  |  |  |  |  |  |  |
| Synonymous | 10,209 | 1.34 | 10,214 | 1.36 | 10,182 | 1.34 | 10,211 | 1.34 | 10,184 | 1.34 |
| Missense | 8,915 | 1.36 | 8,938 | 1.38 | 8,918 | 1.36 | 8,910 | 1.35 | 8,911 | 1.36 |
| Stoploss | 10 | 1.49 | 10 | 1.50 | 10 | 1.53 | 10 | 1.49 | 10 | 1.43 |
| Stopgain | 70 | 2.46 | 70 | 2.48 | 67 | 2.24 | 70 | 2.46 | 70 | 2.45 |
| Frameshift insertion | 24 | 2.22 | 25 | 2.24 | 25 | 2.30 | 24 | 2.21 | 24 | 2.21 |
| Frameshift deletion | 33 | 4.9 | 33 | 5.01 | 33 | 6.08 | 33 | 4.85 | 32 | 4.51 |
| Non-frameshift insertion | 25 | 1.88 | 26 | 1.92 | 26 | 2.01 | 25 | 1.87 | 26 | 1.87 |
| Non-frameshift deletion | 44 | 4.56 | 45 | 4.74 | 44 | 5.20 | 43 | 4.46 | 43 | 4.91 |
| **Disease-associated** |  |  |  |  |  |  |  |  |  |  |
| SIFT:deleterious | 1,623 | 2.86 | 1,625 | 2.90 | 1,627 | 2.82 | 1,623 | 2.85 | 1,622 | 2.85 |
| Polyphen2:probably damaging | 956 | 3.32 | 958 | 3.38 | 958 | 3.25 | 955 | 3.31 | 951 | 3.22 |
| Polyphen2:possibly damaging | 758 | 2.92 | 761 | 2.96 | 758 | 2.88 | 757 | 2.92 | 756 | 2.91 |
| MutationTaster:disease causing | 8,591 | 1.24 | 8,617 | 1.26 | 8,593 | 1.24 | 8,585 | 1.24 | 8,575 | 1.24 |
| ClinVar:Pathogenic | 11 | 1.9 | 11 | 1.75 | 11 | 2.58 | 11 | 1.90 | 11 | 2.08 |

10bp from exon-intron boundary as splice variants

The homozygous pathogenic variants are responsible for recessive Mendelian disorders. To measure the prevalence of pathogenic variants in a healthy individual, we annotated the variants by ClinVar (Landrum et al., 2014). In total, we identified 757 pathogenic variants in all the healthy population, and in average each individual carried 11 pathogenic variants (het/hom = 2.08) (Tables 1 and Table S4). Each genome carried 3.6 ± 2.1 (mean ± SD) pathogenic homozygote variants in Han Chinese population. Additionally, we found that 19 pathogenic variants existed in all four sub-regions (North, Central, South and Lingnan population) were mainly relevant to the immune system, metabolic, and hearing impairment diseases, and 16 out of the 19 (84.2%) variants had a higher frequency (MAF > 0.01) in the Han Chinese population (Table S4), and lower frequency (MAF < 0.001) in 1KG (Genomes Project et al., 2015) and gnomAD database (Lek et al., 2016).

### Genetic Evidence Supported the Geographical Boundaries of the Qinling-Huaihe Line and Nanling Mountains

To explore the Chinese population structure, we performed principal component analysis (PCA) on 2,056 Han Chinese individuals and 205 minority individuals from 29 of 34 administrative divisions of China (Figure 2A). PC1 and PC2 revealed the main genetic structure of the Chinese population, with PC1 displaying a significant population stratification along the north-south cline, reflecting the geographical locations (Figure 2B). The genetic difference of the Han population corresponded to the geographical boundaries of the Qinling-Huaihe River Line and Nanling Mountains. Based on the PCA analysis, the Han Chinese could be classified into four clusters: North Han (Gansu, Hebei, Heilongjiang, Henan, Inner Mongolia, Jilin, Liaoning, Ningxia, Qinghai, Shaanxi, Shandong, Shanxi and Tianjin) (Figure 2B and Figure S8), Central Han (Anhui and Jiangsu) (Figure 2B and Figure S9), where Central Han were closed to North, but embedded in both North and South Han, South Han (Chongqing, Fujian, Guizhou, Hubei, Hunan, Jiangxi, Sichuan, Yunnan and Zhejiang) (Figures 2B and S10), and Lingnan Han (Guangxi, Guangdong and Hainan) (Figures 2B and S11). When the 104 JPT (Japanese in Tokyo, Japan) and 99 KHV (Kinh in Ho Chi Minh City, Vietnam) samples from the 1000 Genomes Projects (1KG) were included, the KHV population formed a cluster overlapping with Lingnan Han, while the JPT population was closer to the North Han Chinese (Figure S12).

**Figure 2.**
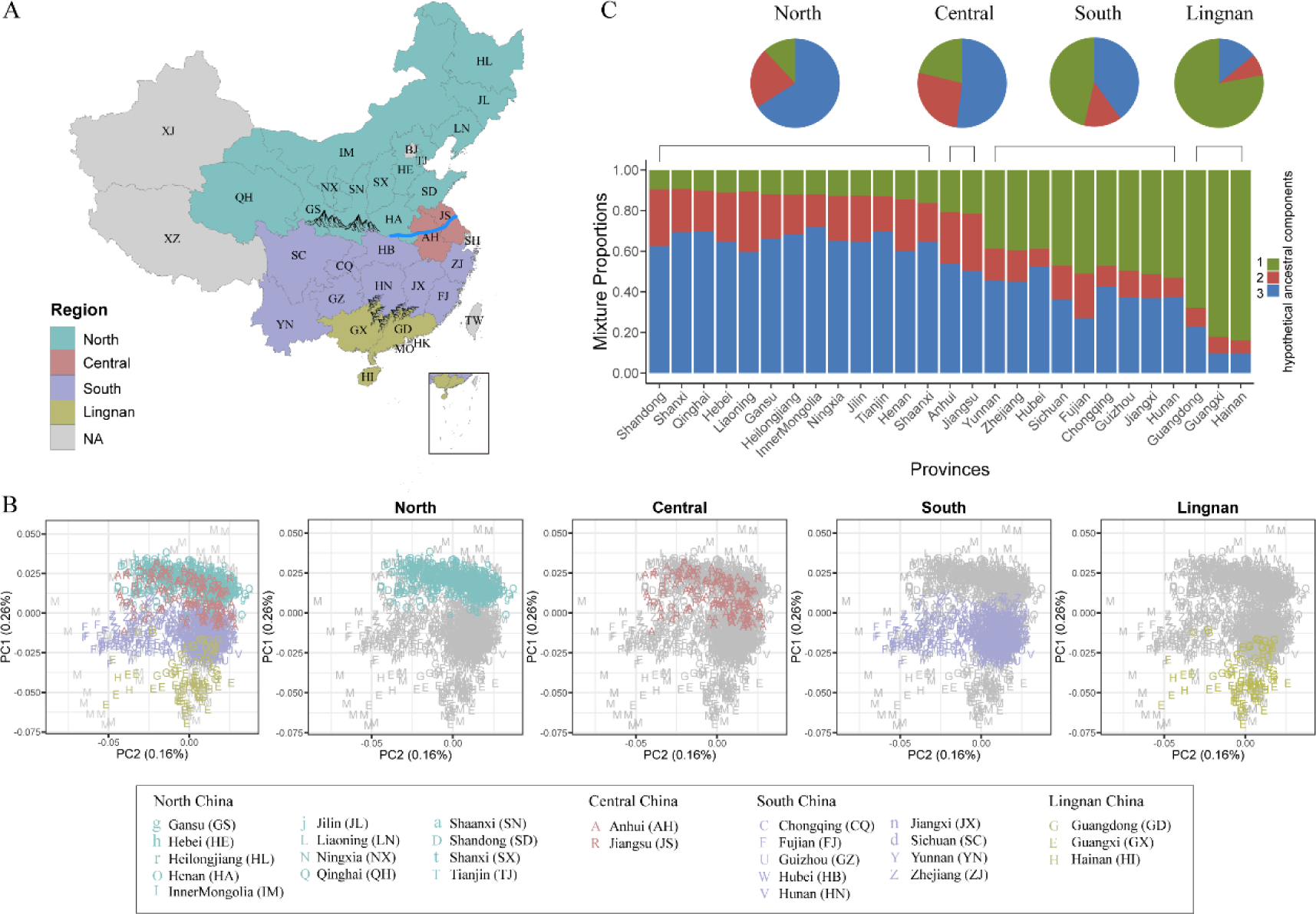
PCA and ADMIXTURE analysis of the Han Chinese populations. (A) A map of the People’s Republic of China showing its 34 administrative divisions. “NA” indicates that the Han Chinese samples were not recruited from that region, which included Beijing (BJ), Hong Kong (HK), Macao (MO), Shanghai (SH), Taiwan (TW), Tibet (XZ), and Xinjiang (XJ). The Qinling-Huaihe River line lies in central China, while the Nanling Mountains are in southern China. (B) Principal Component Analysis (PCA) of the Han and Minority Chinese individuals from four regions. The administrative divisions are shown by the distinct letters. Minority people are marked with “M”. The first two components show the main differentiation between Northern and Southern structures. The Han Chinese populations can be classified into four subgroups: North Han (cyan color), Central Han (dark-red color), South Han (purple color), and Lingnan Han (golden color). (C) ADMIXTURE analysis of 2,056 Han Chinese individuals from 27 administrative divisions for the optimal K value = 3. Each vertical bar represents the average proportion of ancestral components in the regions. The length of each color indicates the percentage of inferred ancestry components from ancestral populations. Provinces are arranged by the value of hypothetical ancestral components 1 in each group. The upper pie charts denote the average proportion of components across individuals from the four subgroups.

We estimated ancestral composition in the Han Chinese population from 27 provinces using the ADMIXTURE program (Patterson et al., 2012). The average number of presumed ancestral populations were calculated in each province with the optimal K = 3. When the value of component 1 was sorted, the four regions were arranged from northern to southern China (Figure 2C). The ancestry fractions of the North Han accounted for about 66% on component 3. The ancestral component of the Central Han was closer to the North Han with 52.1% on component 3, while the admixture components in the South Han were 46.3% on component 1 and 40% on component 3 respectively, which did not show the predominant ancestral components. We found a distinctly higher proportion of component 1 in Lingnan Han, at 78% of ancestry composition compared to other ancestral components.

We also combined the 1KG data (CHB, CHS, CDX, JPT and KHV) and conducted ADMIXTURE analysis from K = 2 to K = 8 to explore the admixture with Asian population (Figure S13). At K = 4, the cross-validation error was the lowest. North Han, South Han, and Lingnan Han showed significantly different clusters, while central Han embodied the ancestral components of both northern and southern populations. In southern China, South Han and Lingnan Han were clearly distinguished from each other, which was consistent with the PCA results. Based on the outcome of the admixture analysis, individuals from the Guangdong province displayed a complex genetic pattern consisting of Lingnan Han and South Han. There were no significant components difference among the Northern provinces. Most components of the CHB population were consistent with North Han, except for part of samples originating from South Han, while the CHS population mainly came from South Han. From K = 2 to K = 6, the JPT population and Han Chinese population were classified into different groups, but more close to the North Han at K = 2. KHV population clustered with Lingnan Han at K = 2, whereas from K = 3 onwards, the KHV population and Lingnan Han clustered separately.

### Population Genetic Structure and Demographic History in Four Sub-regions of the Han Chinese Population

Weir-Cockerham F_ST_ is an allele frequency-based metric to measure the population differentiation due to genetic structure. We calculated pairwise F_ST_ and performed hierarchical clustering for 27 administrative divisions of China and 26 populations of the 1KG. The 27 administrative divisions were mainly clustered into three groups and showed an association with geography (Figure 3A and Table S5). Anhui and Jiangsu provinces, which we designated as Central region of China, were clustered with Northern provinces, indicative of a closer genetic relationship. The other two groups, South and Lingnan, aligned with the regions we designated. Besides, the hierarchical branches suggested that the population differentiation between South and North was smaller than that between the South and Lingnan (Figure 3A), reflecting the relatively shorter genetic distance. The two most remote regions in geography, North and Lingnan, were also found to have the largest population differentiation (Figure 3A). Not surprisingly, the pairwise F_ST_ clustering results between the WBBC and 1KG populations showed that the four designated regions were clustered into the EAS group (Figure S14 and Table S6). In particular, North and Central were clustered with CHB, while South and Lingnan were clustered with CHS (Figure S14). Using the four 1KG continent-level ancestry groups (AFR, EUR, AMR and SAS) as the Non-Chinese population reference, we further investigated the geographical patterns of F_ST_ in 27 administrative divisions of China. The AFR group showed the largest F_ST_ that ranged from 0.14 to 0.15 (Figure S15B and Table S7), indicative of the greatest population differentiation to the WBBC, while the SAS and AMR group yielded the least value (Figure S15A and S15C). The geographical patterns of F_ST_ across the four sub-regions were similar to each other. On average, the Han Chinese in Northern provinces had the relatively closer genetic structure to the Non-Chinese populations of the 1KG. Interestingly, Qinghai province was conspicuously highlighted in the geographic heatmaps, as its pairwise F_ST_ value was obviously smaller than that of other administrative divisions (Figure S15), indicative of the genetic structure particularity of the Qinghai Han Chinese.

**Figure 3.**
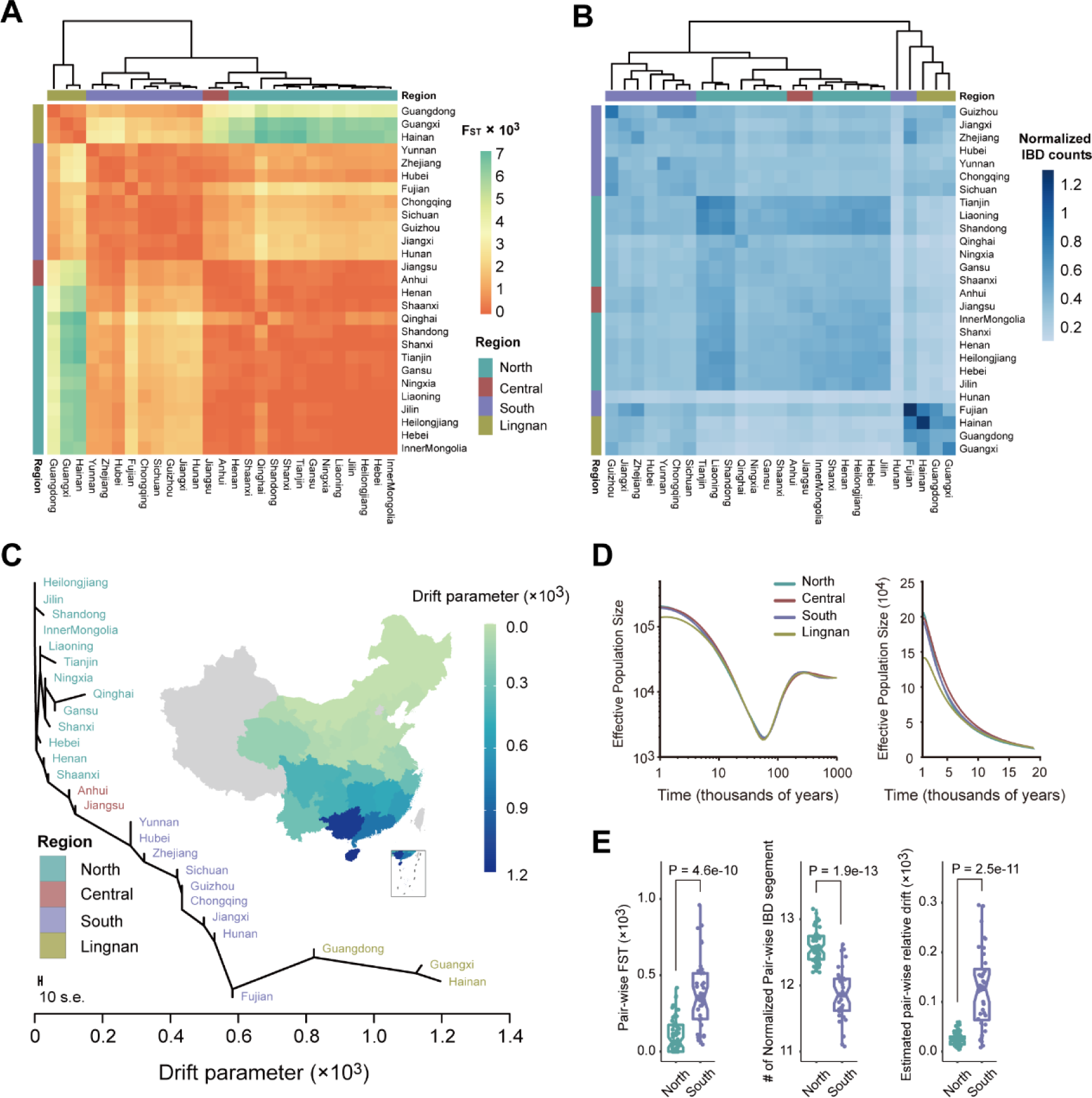
F_ST_, IBD, genetic drift, and effective population size of the Han Chinese populations. (A) A heatmap of pairwise F_ST_ between any two of the 27 administrative divisions in China. The bars on the top and left show the classification of administrative divisions in the four regions. The hierarchical clustering is implemented by the *hclust* function in the pheatmap R package. (B) A heatmap of pairwise IBD segments count between administrative divisions in China. The number of IBD segments is normalized by the sample size of each province (see Methods). The hierarchical clustering is implemented by the same method as the F_ST_ clustering used. (C) A maximum likelihood tree of the Han Chinese in 27 administrative divisions. The plot is rooted in the northernmost province, and the x-axis represents estimated genetic drift. All administrative divisions in the tree are colored by different regions. The scale bar shows ten times the average standard error of the entries in the sample covariance matrix for estimating the drift parameter. The map shows the corresponding geographical distribution. Note that the tree topology for the Northern provinces has a slight difference across the bootstrap replicates due to the extreme homogeneity. (D) Dynamics of effective population sizes of the Han Chinese in four regions. The left panel shows the results on a log–log scale from 1 million to 1,000 years ago and the right panel shows the results on a linear scale over the past 20,000 years. A generation time of 29 years was used to convert coalescent scaling to calendar time. (E) Wilcoxon rank-sum test results for the F_ST_ (left panel), normalized IBD segments (middle panel), and relative genetic drift (right panel) between pairwise Northern provinces and pairwise Southern provinces.

Next, we detected the IBD segments with the logarithm of the odds (LOD) score > 3 across individuals in the WBBC (Lander and Schork, 1994). Unlike the Weir-Cockerham F_ST_, IBD analysis is a haplotype-based approach to reveal the genetic structure and investigate the common ancestry of populations. The total IBD segment counts in each pair of administrative divisions were normalized by the corresponding sample size (see Methods). We then performed the hierarchical clustering based on the matrix of normalized pairwise IBD counts. Similar to the results of F_ST_ clustering, 27 administrative divisions were also mainly clustered into three groups, and individuals from Anhui and Jiangsu provinces were clustered in North (Figures 3A and 3B). Besides, the results showed that most Southern provinces shared more IBD segments with Northern provinces than with Lingnan (Figure 3B), just as observed in the Fst analysis (Figure 3A), suggesting that the Han Chinese in South and North shared more common ancestry than South and Lingnan. The Fujian and Hunan, which we designated as the Southern province, had been found that joined up with Lingnan provinces by the multiple hierarchical branches, indicating that they were more close to Lingnan in ancestry (Figure 3B), and contiguous to Lingnan geographically (Figures 3C and S15).

We inferred the history of effective population size for the Han Chinese, and the results across the four regions were shown in Figure 3D. In the period from 1 million years ago to ∼ 6 thousand years ago (kya), the Han Chinese size histories of four regions experienced almost identical dynamics. From 200 kya to ∼10 kya, the effective population size experienced a steep decline and then grew rapidly, with the lowest point reached at ∼60 kya, which was indicative of a bottleneck, consistent with previous demographic history studies (Genomes Project et al., 2015; Terhorst et al., 2017; Wu et al., 2019). Around 6 kya, the size histories of the Han Chinese from the Lingnan began to deviate from the other three regions, potentially reflecting the existence of a population substructure within the Lingnan Han Chinese (Figure 3D).

Using the Han Chinese in the most northern province (Heilongjiang) of China as the reference, we estimated relative genetic drifts and inferred a rooted maximum likelihood tree between 27 administrative divisions by TreeMix software (Pickrell and Pritchard, 2012). In the result shown in Figure 3C, the relative drift of the provinces and municipalities were in line with the geographic location. To gain a better understanding of the result, we further drew a geographic heatmap that suggested a general genetic drift trend from the North to Lingnan, with the drift parameter increasing as the latitude decreased (Figure 3C). To judge the confidence in the trend and tree topology, we performed ten bootstrap replicates by resampling blocks of SNPs. The trend was repeated in all replicate results (Figure S16). Besides, we found that the tree topology of administrative divisions in Central, South and Lingnan was stable. In the North, however, the tree topology was slightly different across the replicates, indicating that the genetic structures of the Northern administrative divisions were very similar and could not be precisely presented in the tree topology (Figure S16).

Enlightened by the genetic drift estimation results, we further investigated the homogeneity degree in the genetic structure of the Northern and Southern Han Chinese respectively. We performed the Wilcoxon rank-sum tests (Wilcoxin, 1947) for Northern and Southern administrative divisions using their respective pairwise F_ST_ values, normalized IBD segments counts and relative drift parameters. The results showed that the Han Chinese from North had smaller population differentiation (*p*-value = 4.6e-10) and genetic drifts (*p*-value = 2.5e-11), and shared more IBD segments with each other (*p*-value = 1.9e-13) than those from South (Figure 3E). These results suggested that the genetic structure of the Han Chinese in North was significantly homogeneous than those in South.

### Whole Genome-wide Singleton Density Score Analysis

We inferred recent allele frequency changes at SNVs of the Han Chinese population by calculating singleton density score (SDS) using the WGS data. In total, 4,259,171 biallelic SNVs and 17,943,790 singletons from 4,395 Han individuals were conducted for the SDS computation. On chromosome 16p, we found novel significant selection signatures in *SNX29* gene (Figure 4A), which encoded the sorting nexin-29 protein and was ubiquitously expressed in the kidney, lymph node, ovary and thyroid gland tissues (Fagerberg et al., 2014). In *SNX29* gene, more than 30 SNPs showed strong selection signatures (*p* < 5 × 10^-8^), which indicated significant enrichment of selection in this genomic region. Relatively higher DAF was observed on the top SNP rs75431978 (DAF = 0.176, *p* = 1.31 × 10^-15^) in the Han Chinese population, compared to the values obtained in 1000 Genome Project EUR (DAF = 0.003) and AFR (DAF = 0.002) populations. SNX29 was reported to be a biomarker for vasodilator-responsive Pulmonary Arterial Hypertension (Thayer et al., 2020). Although the function of the *SNX29* gene remained unknown, it could be considered a biological target of nature selection pressure in the Han population.

**Figure 4.**
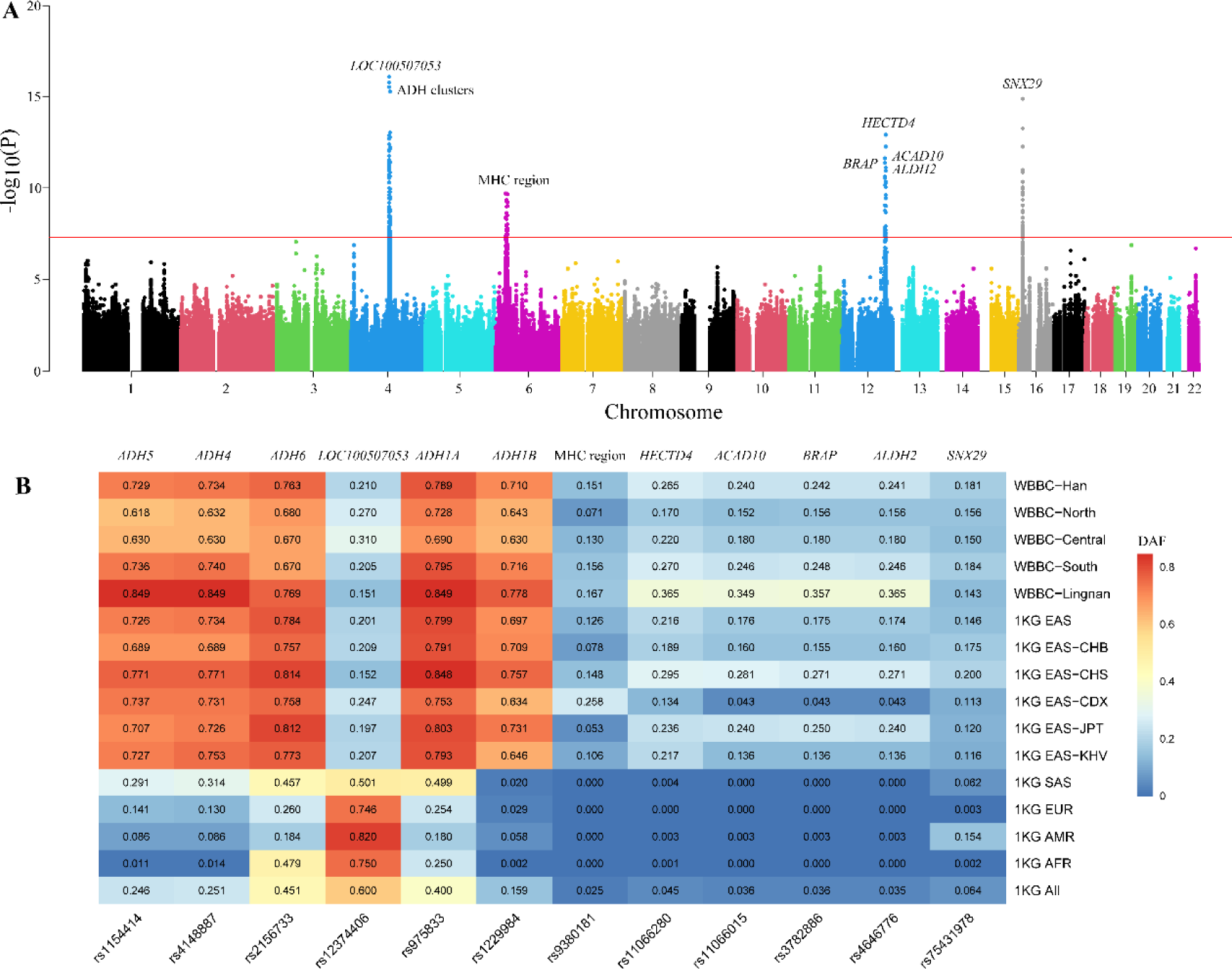
Whole-genome-wide recent selection signatures of the Han Chinese population by singleton density score (SDS) analysis. (A) A Manhattan plot of the natural selection signatures from the WGS data of the Han Chinese individuals. In the x-axis are the 22 autosomes. The y-axis represents the -log10 (*P*) of the two-tailed *p*-values for standardized SDS z-scores. The horizontal red line indicates the significance threshold (p < 5 × 10^-8^). (B) The derived allele frequency (DAF) of SNVs with significant selection signatures in each gene for different populations. The WBBC-Han is all the Han Chinese individuals sequenced by whole-genome sequencing (WGS) in the WBBC cohort. North, Central, South, and Lingnan are the four Han subgroups. EAS, SAS, EUR, AMR and AFR come from the 1000 Genome Project (1KG).

We also confirmed several significant natural selection signals at ADH gene clusters (rs1229984, *p* = 5.51×10^-16^), the MHC region (rs9380181, *p* = 2.04×10^-10^), and BRAP-ALDH2 (rs3782886, *p* = 4.29×10^-12^) (Figure 4A). These three selection signature regions have also been identified in the Japanese population, but the strongest signal loci were at different variants (Okada et al., 2018). The alcohol-metabolizing enzymes such as the alcohol dehydrogenase (ADH) genes, including *ADH1A*, *ADH1B*, *ADH4*, *ADH5*, *ADH6*, and the aldehyde dehydrogenase (*ALDH2*) gene, had an effective impact on the alcohol metabolism pathway and the consequent alcoholism protective effect, which strongly indicated diverse ethnic-specific alcohol consumption patterns (Bierut et al., 2012; Choi et al., 2005; Druesne-Pecollo et al., 2009; Edenberg, 2007; Ehlers et al., 2012). Similarly, the high derived allele frequency (DAF) in this genomic loci, particularly rs1154414 (*ADH5* gene), rs4148887 (*ADH4* gene), rs2156733 (*ADH6* gene), rs975833 (*ADH1A* gene), rs1229984 (*ADH1B* gene), and rs4646776 (*ALDH2* gene) (0.729, 0.734, 0.763, 0.789, 0710 and 0.241, respectively), illustrated corresponding alleles associated with alcoholism in the Han Chinese, when compared with other non-East Asian (Figure 4B and Table S8). Interestingly, we observed a higher-level DAF in these SNVs in South and Lingnan regions compared to the North and Central Han, which reflected the recent regional DAF changes and adaptation in this populous ethnicity and articulated different drinking habits or specific-alcohol consumption. The top SNP rs12374406 in *LOC100507053* gene, as the strongest selection signature on chromosome 4, showed a higher DAF in non-East Asian population, which was also an alcohol dependence risk gene in European-American and African-American populations (Gelernter et al., 2014). In addition, we identified other selection signals in chromosome 12, including rs11066280 (*p* = 1.22×10^-13^) in *HECTD4* gene, rs11066015 (*p* = 2.30×10^-12^) in *ACAD* gene and rs3782886 (*p* = 4.29×10^-12^) in *BRAP* gene, which were limited to EAS ancestry populations. The SNP rs11066280 showed a potential association with blood pressure level (Eom et al., 2017), while rs3782886 may cause a predisposition to an alcohol dependence disorder (Kim et al., 2019). The SNP rs11066015 indicated a strong linkage disequilibrium with rs671 (*p* = 4.59×10^-11^), which was a well-studied variant relevant to alcohol metabolism in East Asian populations (Igarashi et al., 2019). These three genes adjacent to the *ALDH2* gene in chromosome 12q were within a large linkage disequilibrium (LD) block (Levy et al., 2009), revealing that the region had been under positive selection for a long time.

### Signatures of Recent Positive Selection

We employed the iHS test to identify recent natural signatures of positive selective sweeps in the North, Central South, and Lingnan Han populations (Voight et al., 2006). For the iHS test, we computed the fraction of SNVs with |iHS| >2 in the 200 kb non-overlapping genomic windows (Pickrell et al., 2009). The top 1% of genomic clusters were identified as the candidate regions under natural selection. We determined the adaptive candidate genes near SNVs within the 100 kb regions. In total, 130 genomic regions with higher |iHS| scores were found in each population (Table S9-S12). The numbers of overlapping genomic windows of selective sweep regions across the four populations were shown in Figure S17. Only 34 (26%) sweep regions were found in all the four populations. Most regions were shared in two or three of the four subgroups. Averagely, 23.2% of the regions were independent in North, Central and South Han. However, the Lingnan Han had distinctly excess independent sweeps (50, 38.5%), which might be inherited from separate ancestral components, consistent with the conclusion from our demographic history analysis. Importantly, we found the *EDAR* gene in the first three sweep regions in all four subgroups, which have showed the strong signatures of positive selection in East Asians (Mou et al., 2008; Riddell et al., 2020; Tan et al., 2013).

In addition, we conducted Gene Ontology (GO) and KEGG pathway analysis for candidate genes in the top 1% genomic regions with signals of recent selection (Tables 2 and S13). The terms were selected according to *p*-value (< 0.05). The results of the GO analysis showed a significant enrichment of positively selected genes for ethanol metabolic process and ethanol oxidation in four sub-regions, consistent with the selective signatures by whole genome-wide singleton density score (SDS) analysis in the Han Chinese population. We also observed intriguing enrichment of keratinocyte differentiation, epidermal cell differentiation and skin development (*CTSL*, *COL7A1*, *PKP3*, *SEC24B*, *SLITRK6*, *WNT10A*, *KRT10*, *KRT12*, *KRT20* and *KRT23*) in the South and Lingnan Han, which were not present in the North and Central Han populations. KEGG analysis found that 23 pathways were enriched in the Han population with adjusted *p*-values < 0.05 (Table S13). Of these pathways, Southern individuals displayed significantly enriched terms more than the northern population. Tyrosine metabolism and retinol metabolism and fatty acid degradation were identified in four sub-regions.

**Table 2.** Gene Ontology (GO) analysis of candidate genes in top 1% genomic regions with positive selection by iHS.

| GO term | Categories | North | Central | South | Lingnan |
| --- | --- | --- | --- | --- | --- |
| **GO biological process** |  |  |  |  |  |
| GO:0006067 | ethanol metabolic process | 0.0010230 | 0.0154944 | 0.0022526 | 0.0005998 |
| GO:0006069 | ethanol oxidation | 0.0008038 | 0.0154944 | 0.0018229 | 0.0000114 |
| GO:0008544 | epidermis development | - | - | 0.0000594 | 0.0008144 |
| GO:0009913 | epidermal cell differentiation | - | - | 0.0000091 | 0.0003389 |
| GO:0016999 | antibiotic metabolic process | - | 0.0154944 | 0.0008874 | - |
| GO:0017000 | antibiotic biosynthetic process | - | 0.0045203 | 0.0001621 | - |
| GO:0030216 | keratinocyte differentiation | - | - | 0.0000079 | 0.0000755 |
| GO:0031424 | keratinization | - | - | 0.0000002 | 0.0000073 |
| GO:0043588 | skin development | - | - | 0.0000790 | 0.0001520 |
| GO:0050665 | hydrogen peroxide biosynthetic process | - | 0.0096893 | 0.0007227 | - |
| **GO cellular component** |  |  |  |  |  |
| GO:0005882 | intermediate filament | - | - | 0.0000001 | 0.0000004 |
| GO:0045095 | keratin filament | - | - | 0.0000079 | 0.0000755 |
| GO:0045111 | intermediate filament cytoskeleton | - | - | 0.0000002 | 0.0000003 |

Benjamini and Hochberg (BH) correction was applied to enrichment *p* values for multiple testing.
“-” represents that the adjusted *p*-value is >0.05.

### Imputation in the Chinese Population

We evaluated the genotype imputation accuracy of the WBBC, 1KG (Phase 3, v5a) (Genomes Project et al., 2015), CONVERGE (CONVERGE, 2015), and two combined reference panels (WBBC+EAS and WBBC+1KG) in the Chinese population (Figure S18). For the comparison purpose, we extracted 729,958 imputed variants that were shared by the five panels. The variants were then grouped into nine MAF bins (< 0.1%, 0.1%-0.2%, 0.2%-0.3%, 0.3%-0.5%, 0.5%-1%, 1%-2%, 2%-5%, 5%-20% and 20%-50%) to differentiate the detailed imputation performance for variants with different MAF, especially for low-frequency and rare variants, which are usually difficult to impute accurately (Das et al., 2018). We obtained average R-square values (Rsq) from Minimac4 info files (Das et al., 2016) and counted the well-imputed variants (Rsq ≥ 0.8) in each MAF bin. The results showed that the WBBC panel, with almost fifteen-fold more Chinese samples than the 1KG Project, yielded substantial improvement for imputation for low-frequency and rare variants (Figure 5A). The two combined panels, WBBC+EAS and WBBC+1KG, almost tied and possessed both the highest Rsq and well-imputed variant counts for variants with a MAF range of 0.2% to 50%, followed by the WBBC, 1KG and CONVERGE (Figure 5A). For the rare variants with MAF less than 0.2%, WBBC+EAS panel showed the best performance, and the WBBC panel performed roughly the same as the WBBC+1KG (Figure 5A). This result indicated that merging EAS individuals of the 1KG to increase the haplotype size of the WBBC could improve panel’s performance across all MAF bins, but merging the whole 1KG cannot yield more improvement than merging-EAS-only and even not equal to it when the imputed variants were quite rare. Taking all shared variants together, the WBBC+EAS yielded the most well-imputed variants, while the CONVERGE panel imputed the least (Figure 5B). The proportion of imputed variants with Rsq ≥ 0.8 for CONVERGE was the only one under 50% across five panels, even it was population-specific to Chinese (Figure 5C), indicative of the importance of coverage sequencing depth of a reference panel.

**Figure 5.**
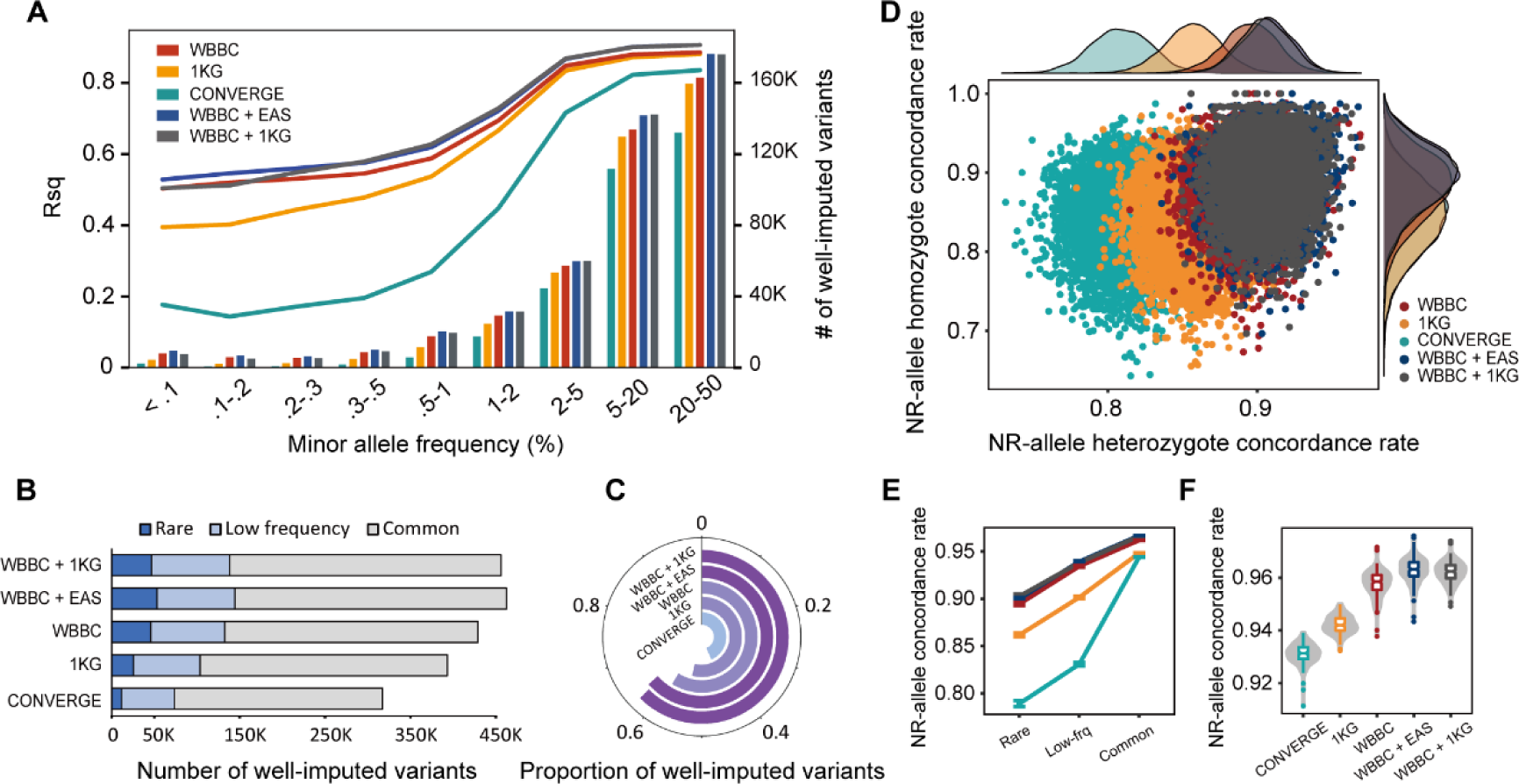
Imputation performance of five reference panels in the Han Chinese. (A) The average R-square (Rsq) and number of well-imputed (Rsq ≥ 0.8) variants in each MAF bin for Chinese imputation by five reference panels. (B) The cumulative number and (C) proportion of well-imputed variants. (D) Non-reference allele (NR-allele) concordance rate distribution (imputed variants vs. array variants). Each dot represents an individual. The x-axis and y-axis denote heterozygote and homozygote, respectively. The plots on the top and right are the corresponding density distributions. The NR-allele genotype concordance rate for (E) rare, low-frequency, and common variants and (F) overall variants (imputed variants vs. WGS variants). The 1KG means 1000G Phase3 and EAS means East Asian group in 1000G Phase 3. All imputations were conducted on chromosome 2.

To comprehensively evaluate the imputation accuracy for the five panels, we further calculated the genotype concordance rate between imputed and genotyped variants by chip array and WGS respectively (imputation vs. chip array and imputation vs. WGS). Since the reference allele was not informative for the evaluation of imputation accuracy, the concordance rate was only calculated in the non-reference (NR) alleles. We randomly masked one-fifteenth sites on QCed chip array for 5,679 individuals, resulting in 2,600 SNPs (chromosome 2 only). The concordance rates of NR alleles were then computed for each sample after imputation. To gain a better understanding of the distribution of the genotype concordance, we separated the NR alleles into homozygote and heterozygote. The result was shown in Figure 5D, with each dot representing an individual and the density plots around the scatter diagram showing the distributions for both NR allele homozygote and heterozygote concordance rates. Two combined panels had the most promising distributions of the concordance rates, which were almost coincident with each other, indicating that the concordance rates for Chinese imputation could barely benefit from the extra population-diverse haplotypes of the reference panel (Figure 5D). Besides, we could know that the peaks of two combined panels in density plots were higher than other panels, indicating that the distributions of concordance rates were more concentrated in the two combined panels (Figure 5D). The performance of the WBBC panel was slightly behind the two combined panels, but was superior to the 1KG and CONVERGE (Figure 5D). We also calculated the NR allele concordance rate between the imputed genotypes and the directly sequenced genotypes. There were 184 samples that were used for both the chip array genotyping and WGS of which 179 met our quality control criteria. All of them had been held-out from the WBBC to avoid the potential artifacts. We divided the variants that shared by the five panels into three variant groups, including rare, low-frequency and common variants. Not surprisingly, the two combined panels performed best and were approximately coincident and very closely followed by the WBBC (Figure 5E). This result suggested that the improvement provided by the EAS and 1KG were unremarkable. Considering all variants together, the WBBC+EAS panel showed the highest NR allele concordance rate, followed by the WBBC+1KG, WBBC, 1KG and CONVERGE (Figure 5F).

Overall, we employed Rsq, and NR allele concordance rate for both WGS and array genotype to measure the imputation accuracy for the five panels. Our results demonstrated the superiority of the WBBC as a reference panel for Chinese population imputation. Compared to the 1KG and CONVERGE, WBBC panel greatly improved the imputation accuracy, especially for the rare and low-frequency variants. Besides, we found that merging EAS haplotypes into the WBBC could improve the imputation accuracy, while the extra diverse haplotypes of the 1KG could barely contribute to it.

### The WBBC Genotype Imputation Server

To facilitate genotype imputation in Chinese population, we developed an imputation server with user-friendly website interface for public use (https://imputationserver.westlake.edu.cn/). Users can register and create imputation jobs freely by uploading their bgzipped array data (VCF-formatted) to our server under a strict policy of data security. To ensure the integrity of array data for next phasing and imputation, some basic QC should be performed, such as removing mismatched SNPs, monomorphism and duplicate SNPs. The server provided a choice of four reference panels to conduct the imputation, including the WBBC, 1KG Phase3, WBBC combined with EAS, and WBBC combined 1KG Phase3. All panels in both GRCh37 and GRCh38 were built to meet different needs. Besides, service of phasing was also provided in our server for users who cannot afford the corresponding heavy computational load. An email of reminder will be sent to the user when the imputation job is finished, and then user can download the imputed genotype data and the corresponding statistics file with an encrypted link. The SHAPEIT v2 (Delaneau et al., 2011) and MINIMAC v4 (Das et al., 2016) were employed in our server for phasing and imputation, respectively. More details including the policy of data security, statistics of four reference panels, and the reference manual were specified in our website.

## Discussion

We initiated the Westlake BioBank for Chinese (WBBC) pilot project and performed the whole genome sequencing at 13.9 × coverage of 4,535 individuals from 29 of 34 administrative divisions of China. We described a comprehensive map of the whole genomic variation in the Chinese population (https://wbbc.westlake.edu.cn) and identified 31.27 (38.6%) million novel variants. Together with 5,841 individuals genotyped by high-density Illumina Asian Screening Array (ASA), we have investigated into the structure of Chinese population, and found that the genetic evidence supported the geographical boundaries of the Qinling-Huaihe Line and Nanling Mountains, which separated the Chinese into four sub-regions (North Han, Central Han, South Han and Lingnan Han). The genetic architecture within North Han was more homogeneous than South Han. We found novel significant selection signatures around *SNX29* gene in the Han Chinese, and confirmed several significant natural selection signals at ADH gene clusters and MHC region. We observed enriched positive selective sweeps of keratinocyte differentiation, epidermal cell differentiation and skin development in the South and Lingnan Han. We provided a comprehensive reference panel for genotype imputation for Chinese and Asian population, and an online imputation server (https://imputationserver.westlake.edu.cn/) is publicly available now for genotype imputation.

The genetic structure of a population defines the level and extent of genetic variation within its constituent subpopulations. Our finding demonstrated that the Han Chinese populations were divided into four sub-regions (North, Central, South and Lingnan), which corresponds to the geographical boundary, the Qinling-Huaihe Line and Nanling Mountains (Five Ridges). Our data did not support the classification of seven subgroups in the Han Chinese as reported previously (Cao et al., 2020). Shuhua Xu et al showed that the Han Chinese was distinguished with three clusters corresponding roughly to northern Han, central Han and southern Han (Xu et al., 2009). Notably, the administrative divisions of North Han and Central Han by Xu et al were consistent with our results, however, the southern Han would be accurately separated into South Han and Lingnan Han by the geographical barrier of the Nanling Mountains and Yunnan-Guizhou Plateau, which had been confirmed by our PCA and ADMIXTURE results. Additionally, the genetic architecture within North Han were distinctly homogeneous, while the ancestral components of admixture in South Han were more diverse. Due to the absence of Han samples in seven administrative divisions (Beijing, Shanghai, Tibet, Xinjiang, Taiwan, Hong Kong and Macao), we have not inferred the population structure in these areas.

Modern Han Chinese have a genetic imprint of early migration patterns and geographical segregation. In ancient times, geographical barriers and inconvenient transportation blocked human migrations. As a result, population subgroups inevitably adapted to local environments in a long period of time, leading to population diversity. Advantageous biological traits were more likely to be passed on to the next generation, influencing the positive selection of ancestral and derived alleles within a population. Adaptive evolution would remove deleterious mutations and increased the beneficial allele frequencies within the gene pool, owing to linkage disequilibrium (LD). The selective pressures acting on the population could result in favorable local adaptive traits. In our study, we identified a novel significant selection signature around *SNX29* gene in the Han Chinese and confirmed several positive selection signatures using the singleton density-based and haplotype-based methods. The ethanol oxidation was the most significant enrichment in the Han Chinese population, which included the ADH gene cluster and ALDH gene cluster. The two shared selective events were maintained by positive selective pressures during Asian population evolution, resulting in a variation in genotype prevalence of these genes in different Asian ethnic groups (Eng et al., 2007). The major histocompatibility complex (MHC) genes carried the significant selection signatures for adaptive autoimmune and infectious diseases in all populations shaped by natural selection over a long time (Meyer and Thomson, 2001). The ectodysplasin A1 receptor (*EDAR*) encodes a member of the tumor necrosis factor receptor family, which was involved in the development of hair, teeth and glands (Fujimoto et al., 2008a; Fujimoto et al., 2008b; Schmidt-Ullrich et al., 2001). The SNV rs3827760 (NM_022336: c.1109T>C p.Val370Val) in *EDAR* gene had been under strong natural selection in the Han Chinese population with a very high allele frequency (0.93). This non-synonymous SNV displayed a higher |iHS| score in four groups (North: 3.08, Central: 3.15, South: 3.15 and Lingnan: 3.65), which indicated a strongest signal of positive selection in this genomic region.

In addition, epidermis is the outermost layer of the skin, which protects the body against pathogens and ultraviolet radiation, and is under adaptive pressure from sunlight duration and intensity. The Qinling-Huaihe line is a geographical dividing line between northern China and southern China, which is the boundary between semi-humid warm temperate continental monsoon climate and humid subtropical monsoon climate in China. Most areas of northern China are dry and cold in winter, whereas there are mild in winter and hot and muggy in summer in southern provinces. The enrichment differences of candidate genes on skin development related traits between northern and southern Han Chinese population might be the results of adaptive pressures selection, including the effects of geography, climate and human migration.

Finally, using R-square and NR-allele concordance rate metrics, we evaluated and compared the genotype imputation performance of the WBBC pilot with two existing panels, the 1KG Phase3 and CONVERGE. Besides, given that the haplotype size of a panel and the genetic background between the panel and array are two crucial factors for imputation accuracy (Bai et al., 2019; Das et al., 2018), we built and evaluated two more combined panels that merged the WBBC with the 1KG and EAS group by the reciprocal imputation approach (Huang et al., 2015). The 1KG Project, which consisted of 2,504 individuals from 26 worldwide populations, is the most diverse and commonly used panel for genotype imputation due to its high quality (Genomes Project et al., 2015). The CONVERGE is the largest and population-specific reference panel for Chinese imputation so far. The quality of variants, however, is not very reliable because of the low-coverage sequencing depth (CONVERGE, 2015). In our study, the WBBC panel yielded substantial improvement for imputation accuracy for low-frequency and rare variants than these two existing panels. The WBBC+EAS and WBBC+1KG panels performed better than WBBC panel alone, and the WBBC+EAS panel yielded highest imputation accuracy for rare variants, the most well-imputed variants and the highest proportion of well-imputed variants. This observation was consistent with and further expanded our previous finding that population-specificity between reference panel and the imputed array was reasonably rigorous for the Han Chinese genotype imputation, and the accuracy benefited from the increasing of haplotype size via extra diverse individuals was limited, especially for rare variants (Bai et al., 2019). Here, to maximize utilization of the WBBC pilot, we provided a large population-specific Genotype Imputation Server, which included the WBBC, 1KG and the two combined reference panels for Chinese sample imputation.

In summary, we characterized large-scale genomic variations in Chinese population. Our finding provided the comprehensive genetic evidence for the geographical boundaries of the Qinling-Huaihe line and Nanling Mountains to divide the Han Chinese population into four subgroups. We elucidated the regional genetic structure and signatures of recent positive selection differences among the Han Chinese ethnic. We also created a user-friendly website and high-performance genotype imputation server for Asian samples. The online resource would practically be important for the genomic variants filtration of monogenic diseases and consequent association with complex traits in the population genetics field.

## AUTHOR CONTRIBUTIONS

H.-F.Z. conceptualized and designed the study. P.C. and W.-Y.B. conducted analysis. S.Y., W.Z. and J.L. conducted the whole sequencing experiments. B.T. and J.L. provided the whole sequencing data from Hunan province. X.Z., P.Z., J.X., M.Q., G.T., S.X., P.G., J.T., M.Y., Y.Q., M.Y. and K.L. contributed to the collection of study samples. Y.L., W.-Y.B., S.G., P.C. and N.L. designed the online website resource. H.-F.Z., P.C. and W.-Y.B. drafted the manuscript, H.-F.Z., B.T., J.L. and S.K. reviewed and edited manuscript. All authors contributed, discussed and approved manuscript.

## Methods

### Samples

The WBBC pilot has enrolled 14,726 individuals with diverse traits across 29 of 34 administrative divisions in China (Provinces, Municipalities and Special Administrative Regions), following the regulations of the Human Genetic Resources Administration of China (HGRAC). A total of 4,535 individuals were whole genome sequenced and 5,841 individuals were genotyped by high-density Illumina Asian Screening Array (ASA) with 73.9 million variants (Table S1). All the participants signed the consent forms. The research program was approved by the Institutional Review Board of the Westlake University.

### Whole genome sequencing and variants calling

Genomic DNA was extracted from peripheral blood samples collected from all the participants using the blood DNA extraction kit (TianGen Biotech, China). We performed the whole genome sequencing on Illumina NovaSeq 6000 system (150 bp paired-end reads) following the standard Illumina library construction and instructions at the KingMed Diagnostics Co. Ltd. The target depth was ∼13× per individual, with about 40 Gb sequencing data.

Variants calling were conducted on all the samples via BWA version 0.7.17 (Li and Durbin, 2010) and GATK4 version 4.1.4.0 (Van der Auwera et al., 2013) (Figure S2). The clean reads in each lane were aligned to the GRCh38 and GRCh37 human reference genome via the BWA mem tool to produce the SAM files, respectively. We used SAMtools (v1.7) view tool to convert SAM format files into BAM files (Li, 2011) and GATK4 MergeSamFiles tool to sort and merge multiples lanes data into one bam. The MarkDuplicates was used to mark the PCR duplicates with a REMOVE_DUPLICATES parameter setting of false. BaseRecalibrator generated the recalibration table by the known sites dbSNP and 1000G VCF resources. ApplyBQSR outputted the recalibrated final BAM files for HaplotypeCaller. We first obtained the GVCF file for each sample and combined all the GVCF files into a single VCF files using GATK4 GenomicsDBImport and GenotypeGVCFs following the suggested pipelines. VariantFiltration was used for filtering excessively heterozygous variants (ExcessHet >54.69) marked with ExcessHet. We calculated the VQSLOD value of each SNV and INDEL variant by setting the max-Gaussians value to 5 and annotating with ReadPosRankSum, MQRankSum, DP, QD, FS and SOR for SNVs, with ReadPosRankSum, DP, QD and FS for INDELs. In the ApplyVQSR filtration, 99.0 was applied for the truth sensitivity level for INDELs and 99.6 for SNPs. All the passed variants were retained for the downstream analyses.

For the X chromosome, we called the genotype in the pseudo-autosomal region (PAR) and non-pseudo-autosomal region (non-PAR), separately. We defined the parameter ploidy with 2 for females and 1 for males in non-PAR, while the parameter was 2 for all individuals in PAR. Then we merged the data together. We only called the genotypes on the Y chromosome for males as the haploid chromosome.

### Sample and variant filtrations

The sex estimation and confirmation were analyzed by the ratio of sequencing depths aligned to the X chromosome and autosomes (Wu et al., 2019). The inferred sex was consistent with the self-reported sex for each sample. The FREEMIX scores were used to estimate DNA contamination by verifyBamID version1.1.3 with –maxDepth 100 --precise --minMapQ 20 --minQ 20 --maxQ 100 and the allele frequencies inferred from our genotyped data (Jun et al., 2012). In total, 15 samples with FREEMIX scores > 0.05 were excluded (Figure S19). We identified the duplicates samples by KING version 2.2.4 --duplicate and removed 31 duplicated individuals or MZ twins (Manichaikul et al., 2010). Finally, 4,489 samples were retained in the final cohort.

In the raw calling set (GRCh38 build), 99,958,705 SNVs and INDELs on autosomes, 4,194,725 SNVs and INDELs on sex chromosomes were identified. Of these, 1,067,527 variants were excluded by ExcessHet. At a truth sensitivity of 99.6% for SNVs and 99% for INDELs, 11,183,295 SNVs and 2,694,079 INDELs were removed. We used the bcftools filter to remove the 2,938,593 SNVs and 2,645,756 INDELs variants closer to INDEL with SnpGap 3 and IndelGap 5. Then we filtered 10,462 INDELs with a length more than 50 bp. Finally, we excluded 1,953,082 variants with a HWE *p* value < 10^-6^ by VCFtools (v 0.1.13) (Danecek et al., 2011).

### Variant annotations

The functional annotation of variants were performed with the ANNOVAR tool (Wang et al., 2010). We annotated the gene name, protein change, location and function for all the variants. The pathogenic or benign of variants were annotated by SIFT (Danecek et al., 2011), PolyPhen-2 (Adzhubei et al., 2013), MutationTaster (Schwarz et al., 2014) and ClinVar version 20200728 (Landrum et al., 2014).

### Genotyping

The 5,841 samples of the WBBC Project and 184 individuals (13.3×-54.7×) sequenced by WGS were genotyped by ASA-750K (Asian Screening Array) BeadChip designed for the East Asian population. The genotype call rates for each sample were more than 95%. We computed the allele frequencies in the Chinese population using 5,841 samples and 484,554 SNP variants passed the filtrations (--geno 0.05 --hwe 0.000001 and --maf 0.01) by Plink version 1.9 (Chang et al., 2015) and were consequently retained for further analyses.

## QUANTIFICATION AND STATISTICAL ANALYSIS

### Evaluation of genotype concordance

We applied the sequencing and genotype data from 184 individuals to estimate the whole genome sequencing calling accuracy. The genotype from SNP arrays were considered as the reference allele and our calling variants were test set. We also conducted the LD-based genotype refinement for the low confidence genotypes and missing sites via BEAGLE 5.1 with default settings (Browning et al., 2018). We computed the heterozygote disconcordance, non-reference genotype concordance, homozygote genotype concordance, specificity and non-reference sensitivity for the shared variants (Figure S20) (Linderman et al., 2014).

### PCA, ADMIXTURE and effective population size inference

We removed the variants in imputed dataset by Rsq ≤ 0.95, and merged it with our sequencing dataset by GATK v4.1.4.0 (McKenna et al., 2010), resulting in 9,996 individuals and 2,016,533 bi-allelic SNPs. We further merged the WBBC dataset with the 1KG Project. After filtering SNPs by MAF ≤ 0.01, a total of 1,857,766 bi-allelic SNPs with 100% call rate were left for subsequent analysis. We noted that the participants of the WBBC Project mainly came from three provinces of China, including Jiangxi (23.8%), Shandong (26%) and Hunan (31.4%). To avoid the potential bias of oversampling certain provinces (McVean, 2009), we randomly extracted 150 samples from each of the three provinces. Finally, 2,056 Han population individuals, 205 Minority population individuals, and 2,504 1KG individuals were included. We then performed PCA (Menozzi et al., 1978), ADMIXTURE (Alexander et al., 2009) and inference of effective population size. Note that the minority population individuals were held-out from each province group.

We excluded the SNPs with HWE p value < 1×10^-6^, MAF < 0.05 and genotype missing > 0.05 using the Plink software (Chang et al., 2015). Then we performed the linkage disequilibrium based SNP pruning with --indep-pairwise 50 10 0.5. The final data sets had 338,275 bi-allelic SNPs for PCA and ADMIXTURE analyses. We used the smartpca command from the software EIGENSOFT (v6.1.4) (Price et al., 2006) and calculated the components for the first ten PCs. PC1 and PC2 were selected for the genetic diversity comparison, which were plotted by in-house R scripts.

ADMIXTURE analysis were conducted with 2,056 Han individuals, 103 CHB (Han Chinese in Beijing, China), 105 CHS (Han Chinese South, China), 93 CDX (Chinese Dai in Xishuangbanna, China), 104 JPT (Japanese in Tokyo, Japan) and 99 KHV (Kinh in Ho Chi Minh City, Vietnam) individuals from combined dataset by ADMIXTURE version 1.3.0 using default parameters (Patterson et al., 2012). To obtain the optimal K value, we analyzed the ADMIXTURE with 10 random seeds for each K ranging from 2 to 8. The default 5-fold cross-validation procedure was carried out to estimate prediction errors. The K value with the highest log-likelihood was selected as the most probable model. We further estimated the history of effective population size for four regions using SMC++ (Terhorst et al., 2017). Using the ancestral components analyzed by ADMITURE with K = 4, we designated 10 most representative samples with the high sequence-depth as the distinguished lineage sample for each region. We followed the suggestion of SMC++ authors and masked all low-complexity regions of the genome using the 1KG Phase3 supported data (Genomes Project et al., 2015), and kept all left bi-allelic SNPs for next analysis. For each region, we repeated SMC++ 10 times according to each distinguished lineage sample. The combined results were used to form the composite likelihood for the final estimation. The per-generation mutation rate was set at 1.25e-8 and a generation time of 29 years was used to convert coalescent scaling to calendar time (Terhorst et al., 2017; Wu et al., 2019).

### F_ST_ statistics, IBD analysis and genetic drift estimation

We next performed F_ST_ statistics (Weir and Cockerham, 1984), genetic drift estimation and identity-by-descent (IBD) analysis. We calculated weighted Weir-Cockerham F_ST_ estimates for each pair of the WBBC provinces and 1KG populations using VCFtools v0.1.13 (Danecek et al., 2011) based on 1,857,766 bi-allelic SNPs. The window size was set to 50,000 and step size to 5,000. We built F_ST_ values matrix and performed hierarchical clustering with it using complete-linkage method implemented in the *hclust* function in the pheatmap package in R.

The IBD analysis was based on haplotypes of individuals. The genome-wide IBD segments were identified for all pairwise Han Chinese from 27 administrative divisions of China using Refined IBD software (Browning and Browning, 2013) with default settings. We built the IBD counts matrix for each pair of administrative divisions. Given that the sample size of 27 administrative divisions were different, we normalized the total IBD counts by sample size. For the IBD segment counts within administrative divisions (for example, province ‘A’), *IBD_normalized counts of A_ = IBD_total counts of A_ / comb(N_A_)*, where *comb* was the combination function in math and *N_A_* was the sample size of province ‘A’. For the IBD segment counts between two administrative divisions (for example, province ‘A’ and ‘B’), *IBD_normalized counts of A vs. B_ = IBD_total counts of A vs. B_ / N_A_ * N_B_*, where *N_A_* and *N_B_* were the sample size of province ‘A’ and ‘B’ respectively. The hierarchical clustering was then performed based on the matrix by using the same method as F_ST_ clustering.

We computed relative genetic drift estimates between each province using TreeMix v1.13 with default settings on the same SNPs as the F_ST_ analysis used (Pickrell and Pritchard, 2012). The genetic drift was represented by a ‘drift parameter’ in TreeMix, more details were described elsewhere in the study (Pickrell and Pritchard, 2012). A maximum likelihood tree for the Han Chinese population from 27 administrative divisions was then plotted. Note that the Heilongjiang province, which was located in the northern most part of China, was set as the reference point. For judging the confidence in our tree topology, ten bootstrap replicates were generated by setting the -bootstrap -k flag ranging from 10 to 100 (step-size = 10) to resample blocks of contiguous SNPs for drift parameter estimation. Plink version 1.9 was used in this part to calculate allele counts of SNPs for reformatting of input data that the software required (Chang et al., 2015).

### Calculation of the singleton density score

The Singleton Density Score (SDS) can be applied to infer recent allele frequency changes in the past 2,000-3,000 years by calculating the distance between the nearest singletons on either side of a test-SNP using whole-genome sequence data (Field et al., 2016). In our data set, 73.91 million autosomal variants were identified in 4,395 Han Chinese samples, of which 68,492,157 were bi-allelic SNVs in all individuals. We filtered the SNVs by Hardy-Weinberg equilibrium (*p* < 1×10^-6^). We downloaded the Homo sapiens ancestral annotation information from the Ensembl release-98. SNVs without defined ancestral allele were subsequently removed. Additional SNPs were excluded by MAF <5% and less than 5 individuals for each of the three genotypes. The final data set included 4,259,171 SNVs and 17,943,790 singletons for the SDS computation.

Gamma-shape was estimated with Gravel_CHB as a demographic model for each derived allele frequency (DAF) bin by 0.005 from 0.05 to 0.95. The haplotypes was set to 8,790, twice the number of individuals. We excluded the centromeres and heterochromatic regions with chromosome boundaries files. The skip boundary missing singletons fraction threshold was 0.5. The raw SDS scores were computed using recommended scripts and standardized within each 1% bin of DAF for each chromosome by calculating z-scores. Two-tailed p-values were converted by whole genome-wide standardized SDS z-scores.

### Calculation of iHS values

To detect the genomic signatures of recent positive selection, we computed the integrated haplotype score (iHS) using the R package rehh v3.1.0 (Gautier et al., 2017; Voight et al., 2006). The data from 2,860 North, 148 Central, 5,274 South and 92 Lingnan Han Chinese individuals were extracted from the imputed and phased files. In total, 1,967,791, 1,897,093, 1,981,861 and 1,853,882 biallelic SNVs were obtained in all autosome chromosomes in four Han populations respectively. The SNVs were further filtered by Hardy–Weinberg equilibrium (--hwe 0.000001) and minor allele frequency (--maf 0.01) using the Plink software (Chang et al., 2015). The ancestral allele of SNVs were defined by the data downloaded from Ensembl release-98. We removed SNVs without an ancestral allele state.

In total, 1,725,164 SNVs in North population, 1,712,580 SNVs in Central population, 1,720,051 SNVs in South population and 1,685,839 SNVs in Lingnan population passed quality control and were retained for statistical analysis. We performed iHS statistics independently for the population. The absolute values of the iHS scores were taken to analyze the data. We calculated the fraction of SNVs with |iHS| > 2 in 200 kb non-overlapping genomic windows (*N*_|iHS|>2_ / *N*_total_) and filtered the windows with < 20 SNVs (Pickrell et al., 2009). The genes located in the top 1% of windows were considered to be significant regions. The genes or genomic regions were defined within 100 kb of the identified non-overlapping SNVs. We performed the Gene Ontology (GO) and Kyoto Encyclopedia of Genes and Genomes (KEGG) pathway enrichment analysis for the adaptive candidate genes using R package clusterProfiler v3.16.0 (Yu et al., 2012).

### Reference panel construction

The multi-allelic sites were split into bi-allelic sites via the BCFtools norm tool version 1.7 (Li et al., 2009). We filtered the variants with --max-missing 0.9 and --hwe 0.000001 using VCFtools version 0.1.13 (Danecek et al., 2011). In total, 508,196 variants were excluded. BEAGLE version 5.1 was used to perform haplotype phasing of all 4,489 samples with default settings. We conducted the haplotype re-phasing with SHAPEIT version 2 (r900) by windows 0.5, states 200 and effective-size 14,269 (Delaneau et al., 2013). Finally, the SHAPEIT haplotypes were converted into VCF format files.

### Quality control, pre-phasing and imputation

The rigorous variant-level and sample-level quality control were then performed as following steps: we kept autosome bi-allelic SNPs and calculated genetic relationship matrix across all individuals using variants with MAF > 0.01 by GCTA v1.91 (Yang et al., 2011), and then samples with the pairwise genetic relationship coefficient > 0.025 were thought to be cryptically related and removed; the variants and samples with a missing call rate > 5% were excluded by Plink version 1.9 (Chang et al., 2015); the variants deviating from Hardy-Weinberg equilibrium at *p* < 10e-6 or with MAF < 0.01 were also excluded. Finally, 5,679 individuals and 470,279 bi-allelic SNPs on autosomes passed the filters and QC. We pre-phased the array dataset by SHAPEIT v2 setting the effective-size parameter to 14,269 as the software recommended for the Asian population (Delaneau et al., 2011). Imputation was then performed with our own haplotype reference panel, which consisted of 8,978 haplotypes at 34,948,874 SNPs (no singleton), by MINIMAC v4 (Das et al., 2016). The length of chunks was set to 20MB with a 4MB overlap between contiguous chunks for the imputation. We employed R-square (Rsq) to control the quality of imputed results and filtered out variants with Rsq ≤ 0.95.

### Reference panel evaluation for imputation in the Chinese population

We evaluated the accuracy of genotype imputation for five reference panels in the Chinese population. These panels included the most widely used panel, the 1KG (Genomes Project et al., 2015), and the largest Chinese-specific panel CONVERGE (CONVERGE, 2015), and our own WBBC panel, and two combined panels that merged the WBBC datasets with the 1KG and EAS respectively. The imputation accuracy of these panels was then compared with each other by three different metrics. The design for the entire evaluation was detailed in Figure S18.

The 1KG Project reference panel (Phase 3, v5a) was downloaded from the ftp sites (http://ftp.1000genomes.ebi.ac.uk/vol1/ftp/release/), and the CONVERGE Project reference panel was downloaded from the European Variation Archive (http://ftp.ebi.ac.uk/pub/databases/eva/PRJNA289433/). For the 1KG, CONVERGE and WBBC reference panels, we split multi-allelic variants into multiple bi-allelic variants and removed singletons and doubletons (minor allele counts, MAC ≤ 2) by using BCFtools. Besides, there were 184 samples that were included in both the WGS and DNA array genotyping for the evaluation purpose. These samples were held-out from the current WBBC panel. Finally, we obtained 3,284,591 variants and 5,008 haplotypes for the 1KG, 1,115,342 variants and 23,340 haplotypes for the CONVERGE, and 2,089,508 variants and 8,610 haplotypes for the WBBC. Note that all the manipulations were conducted on chromosome 2 (Huang et al., 2009; Wu et al., 2019). For two combined reference panels, the WBBC+1KG and WBBC+EAS, we employed the reciprocal imputation approach to implement the combination to preserve maximal variants (Huang et al., 2015). The EAS dataset was directly extracted from the 1KG, and sites with MAC equals zero were removed subsequently. We reciprocally conducted imputation for the WBBC/1KG and WBBC/EAS, and then respectively excluded 2,663 and 2,142 INDELs with incompatible alleles in panels that could fail the next panel-merging. BCFtools was used to finally merge the reference panels (Li et al., 2009). Eventually, the WBBC+1KG combined panel consisted of 13,618 haplotypes at 4,450,989 variants, with 917,784 variants shared by both panels. The WBBC+EAS combined panel consisted of 9,618 haplotypes at 2,411,382 variants, between them, 849,281 variants were shared. We extracted chromosome 2 from our QCed chip array dataset and randomly masked one fifteenth SNPs (Huang et al., 2009; Wu et al., 2019), a total of 5,679 individuals were included and 2,600 SNPs were masked for the next evaluation.

We transformed the format of five panels into M3VCF and performed genotype imputation by jointly using Minimac3/4 (Das et al., 2016). The length of chunks for imputation was set to 20MB with 4MB overlapped between contiguous chunks. The accuracy of different reference panels was evaluated by three metrics. In the first one, the estimated value of the squared correlation between imputed genotypes and true, unobserved genotypes (i.e., R-square) (Das et al., 2016), was calculated based on the imputed dosage and produced with the imputation results by Minimac4. This value was also the most commonly used metric. In this study, an imputed variant with the Rsq ≥ 0.8 was considered as ‘well-imputed’. The second metric was non-reference allele (NR-allele) concordance. The variants that had been masked in the beginning were imputed by different panels. We then calculated the NR-allele concordance between imputed genotypes and the original ones in chip array for each individual (Imputed vs. Array) (Huang et al., 2009). The third metric was similar to the second, but the NR-allele concordance was calculated between imputed genotypes and WGS genotypes by the samples that we hold-out (Imputed vs. WGS). The definition of concordance and corresponding formula was specified in Figure S20 (Linderman et al., 2014).

### Genotype imputation server

Using the WBBC Phase 1 WGS data and 1KG Phase 3 data (Genomes Project et al., 2015), we developed a genotype imputation server for public use. We included the WBBC and 1KG reference panel in the server and re-constructed two combined panels, the WBBC+EAS and WBBC+1KG. All panels were built in both GRCh37 and GRCh38 version, and singletons were excluded. MINIMAC v3 (Das et al., 2016) was used here to build genotype data in the M3VCF format to save the computational memory. We developed the pipeline in Python and Shell, and employed MySQL for the management of data. For the VCF-formatted array data uploaded by users, validity of data would be checked first. Before the actual imputation, there were some basic filtering steps conducted by BCFtools (Li et al., 2009), including removing all mismatched SNPs, monomorphism, and duplicate SNPs. The 1KG was used here as the allele reference. The next phasing and imputation were performed using SHAPEIT v2 and MINIMAC v4 (Das et al., 2016; Delaneau et al., 2011). We specified a policy of data security to protect the user’s data across the entire interaction process with the server. Also, we wrote a help manual and illustrated all processes of our pipeline to facilitate users. Detailed information could be found in our website (https://wbbc.westlake.edu.cn).

### Data and Code Availability

The allele frequencies of all variants and genotype imputation server are available via the website (https://wbbc.westlake.edu.cn). Raw sequencing data have been deposited to the CNGB Sequence Archive (CNSA) of China National GeneBank (CNGBdb) with accession number (CNP0001516) (https://db.cngb.org/cnsa/). The application forms are required for researchers and the study must conform to the regulations of the Human Genetic Resources Administration of China (HGRAC). Researchers who request access to the raw genetic data must get permission from Ministry of Science and Technology of the People’s Republic of China and the Institutional Review Board of the Westlake University.

---

**Figure S1.**
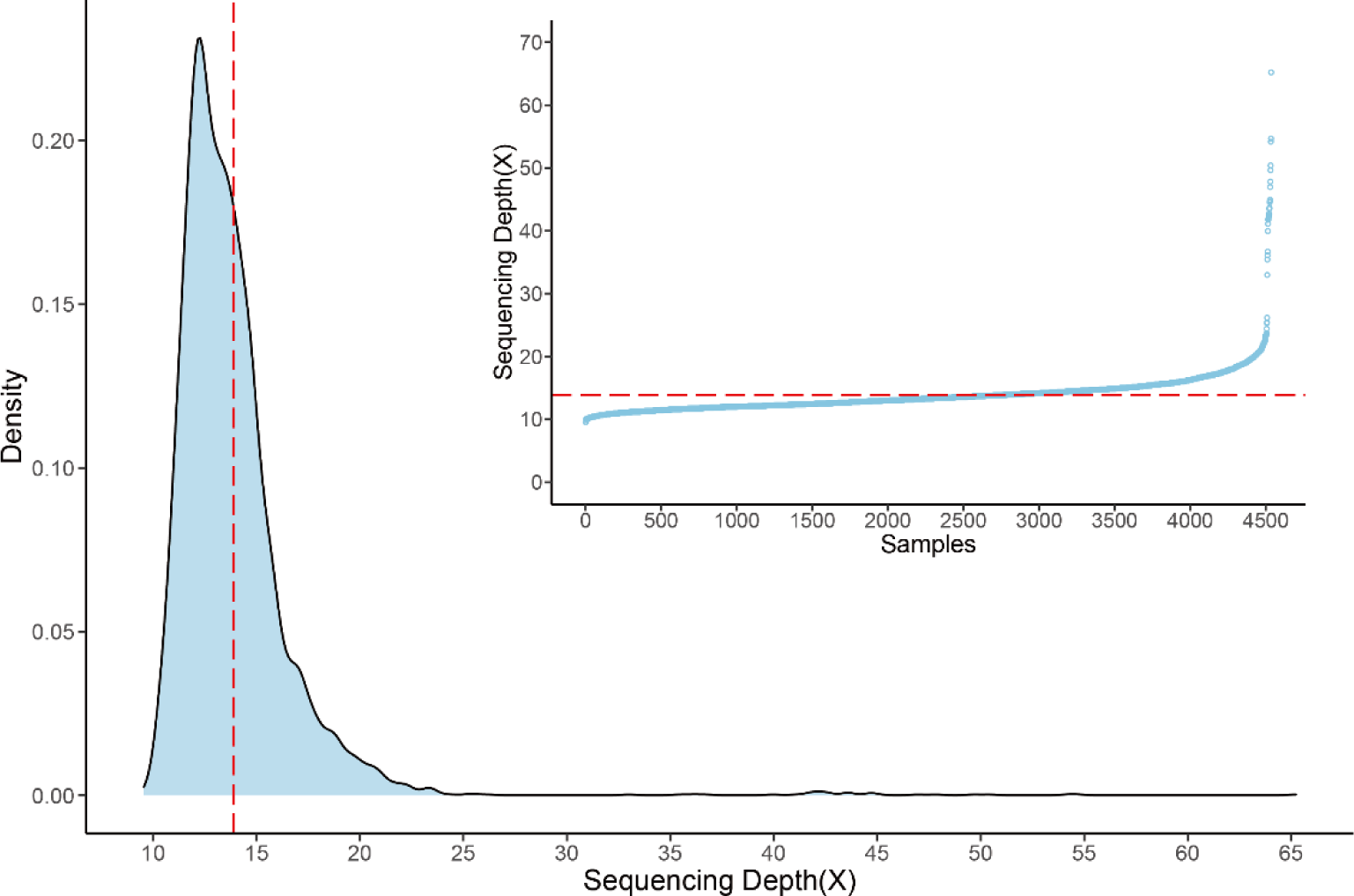
The sequencing depth of all 4,489 samples. The central red dot lines are the median. The inner chart represents the sequencing depth by sorted samples. The area plot indicates the density of sequencing depth.

**Figure S2.**
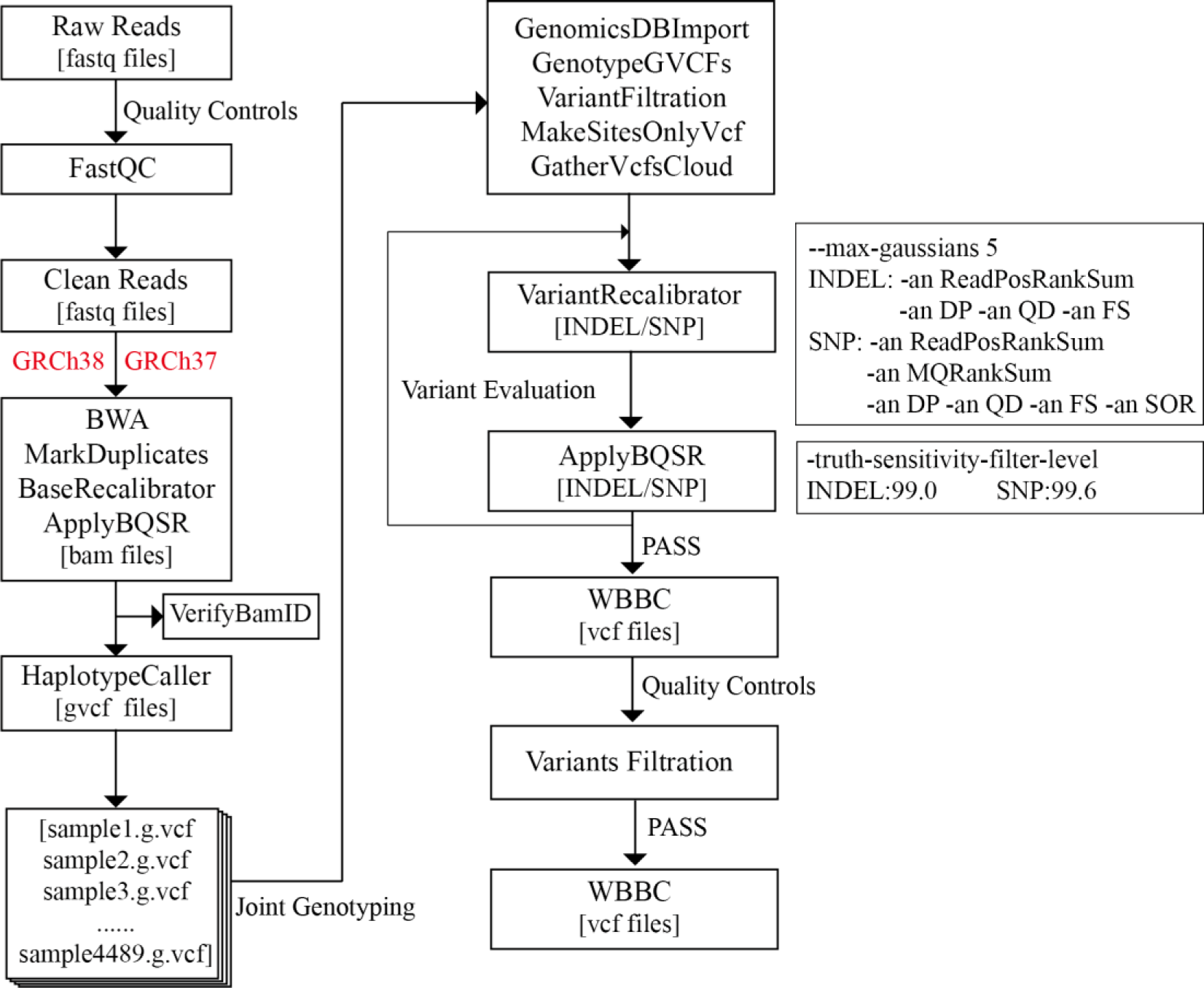
Schematic representation of whole-genome sequencing analysis pipelines. The BWA and GATK4 recommendation pipeline and variant analysis strategy were performed on the WBBC-cohort dataset with the GRCh38 and GRCh37 human reference genome, respectively.

**Figure S3.**
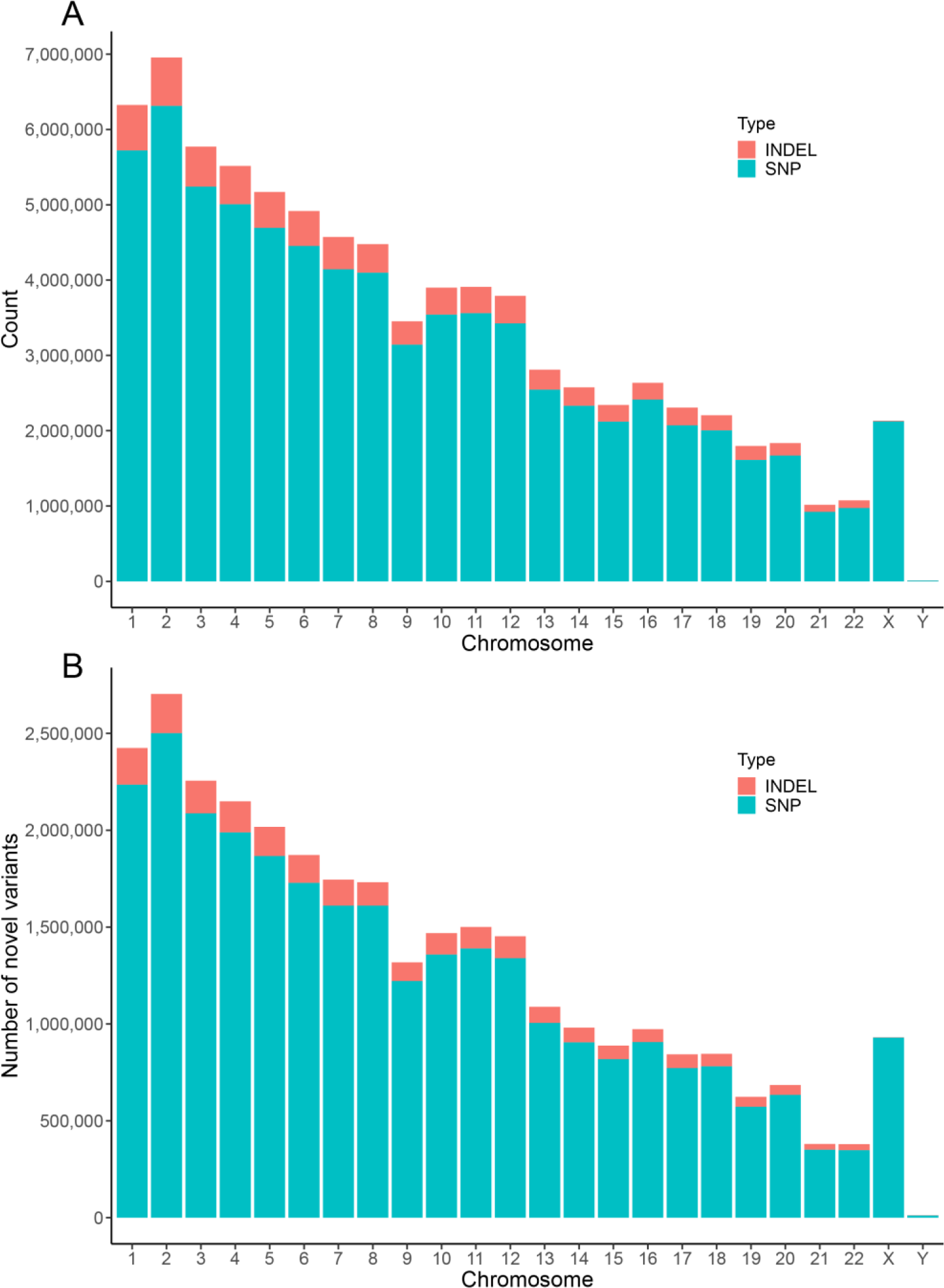
The distribution of variants per chromosome. Green represents the SNVs and red denotes the INDELs. (A) The total number of variants detected in each chromosome. (B) The novel variants based on dbSNP build 151 and their distribution in each chromosome.

**Figure S4.**
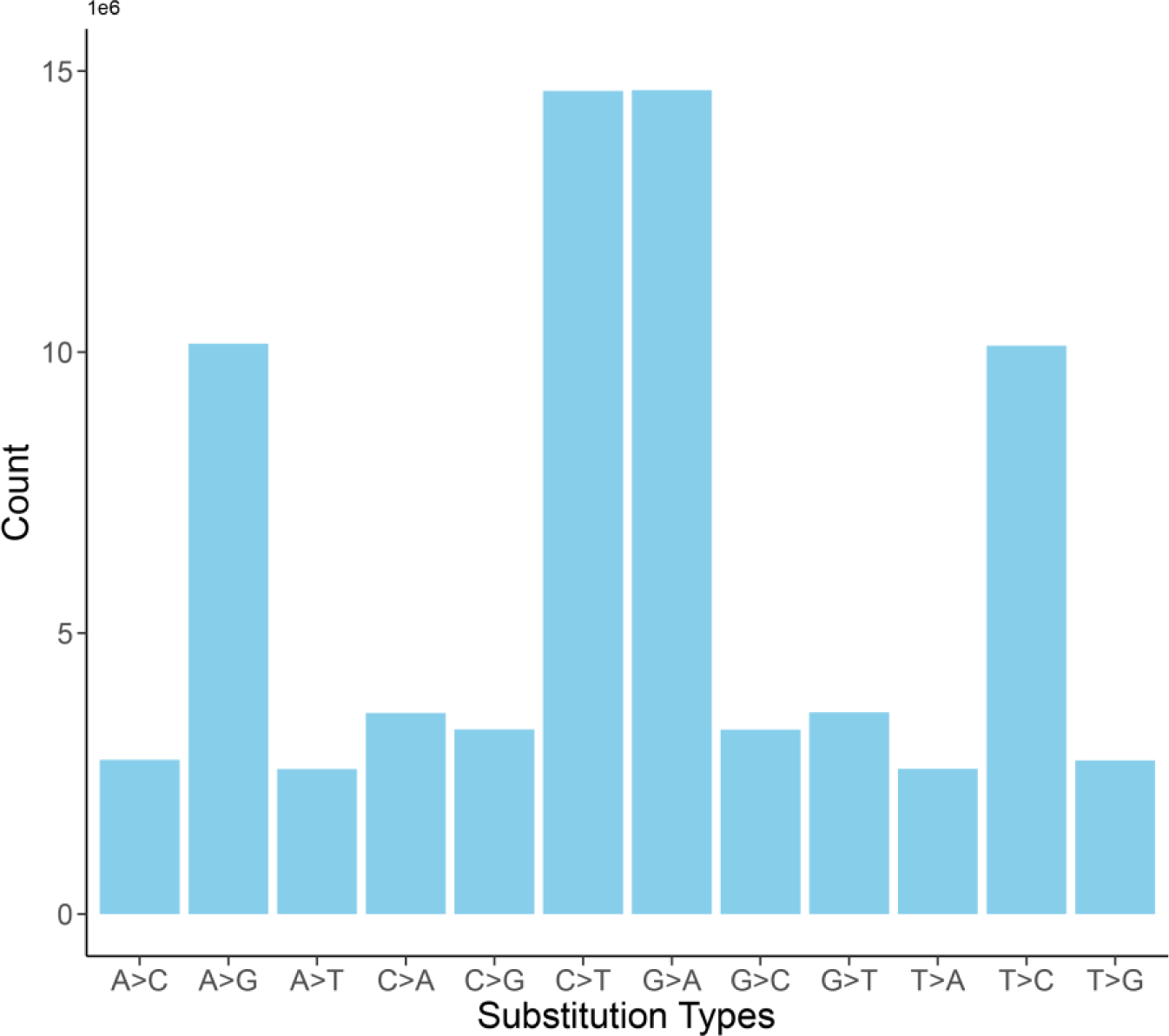
Histogram of the SNVs substitute types in all chromosomes. The x-axis indicated the substitution types. The y-axis represented the number of variants.

**Figure S5.**
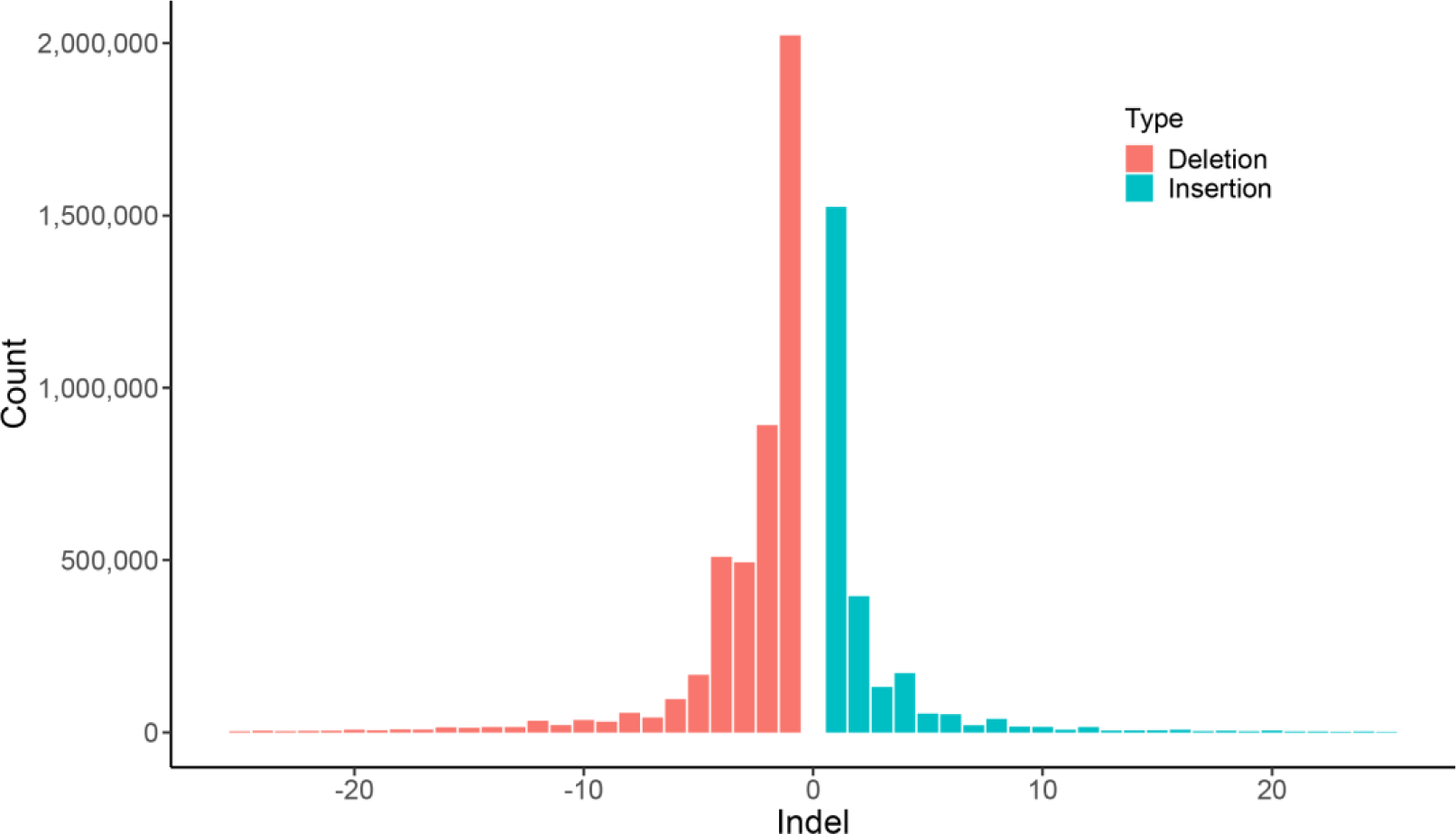
Distribution of the length of INDELs variants. The x-axis was the length of variants from −50 to 50 bp, in which deletion variants were filled with red color and insertion variants with green color. The y-axis represented the number of variants.

**Figure S6.**
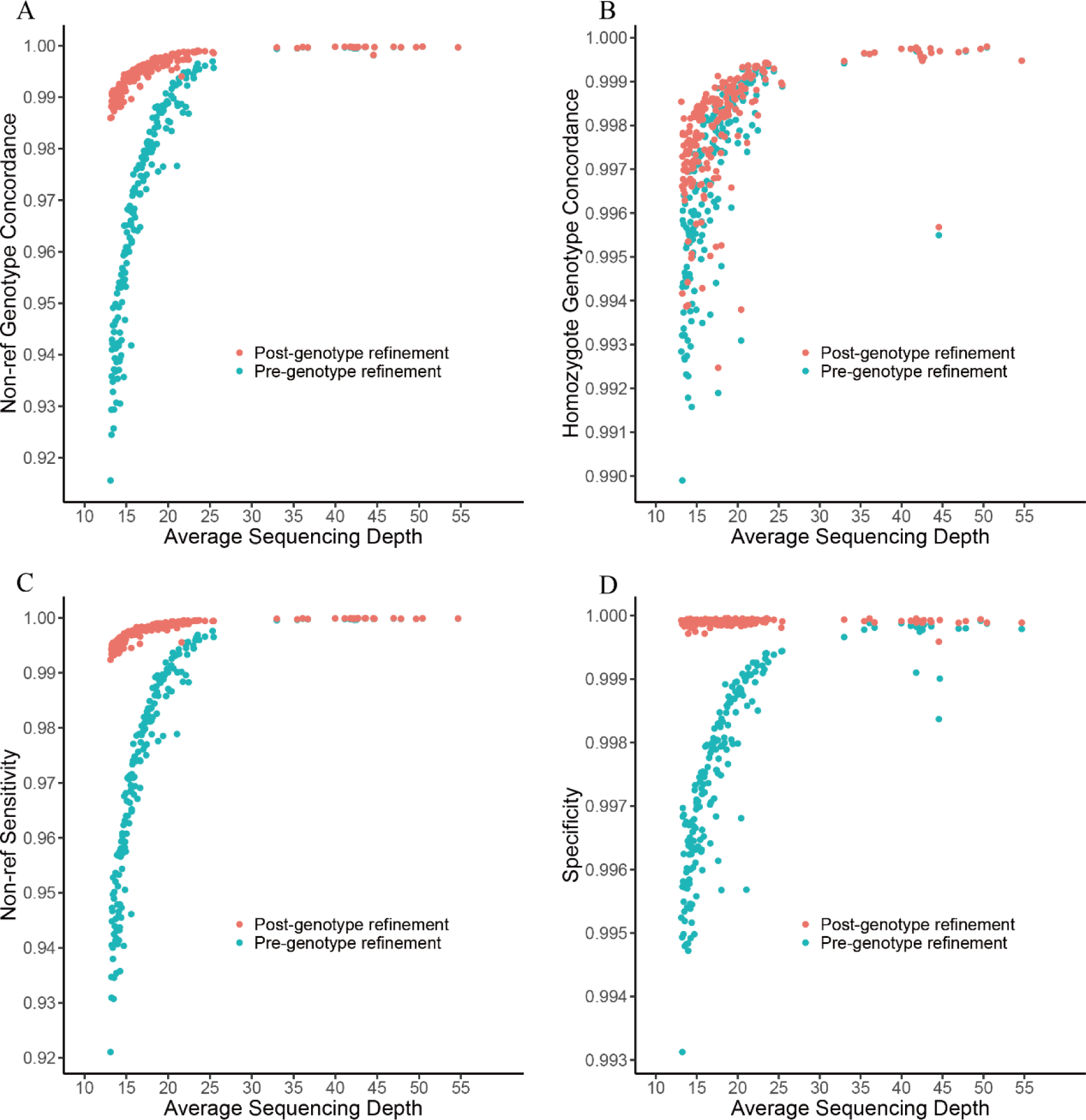
A comparison of the genotype concordance between WGS and SNP array data. (A) Non-reference genotype concordance, (B) homozygote genotype concordance, (C) non-reference sensitivity, and (D) specificity. The LD-based genotype refinement was conducted by BEAGLE software version 5.1. The red dots indicate the average proportion of post-genotype refinement and the green dots denote the raw genotype calls.

**Figure S7.**
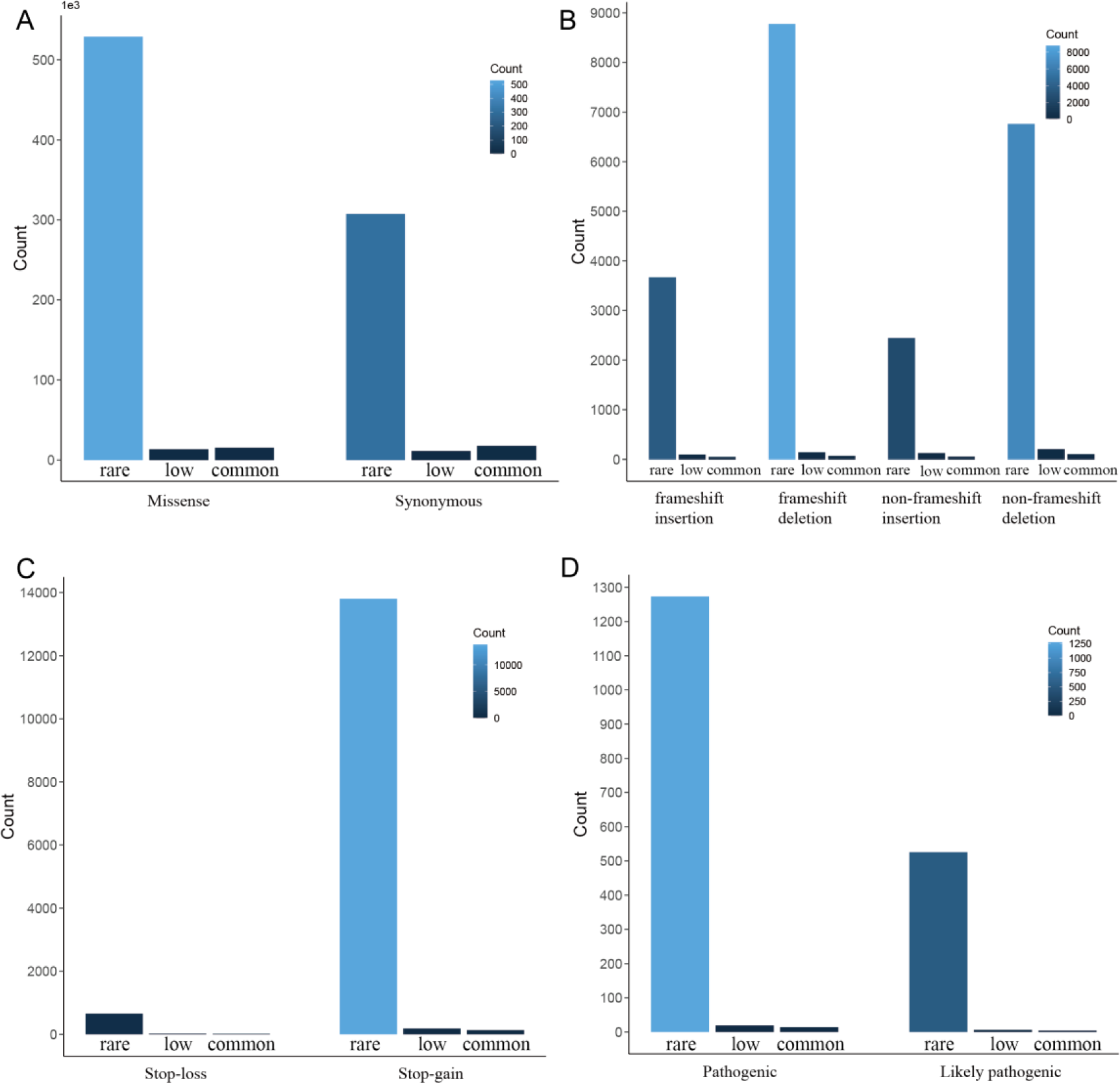
Functional annotation and distribution of variants in three frequency bins. Rare allele (< 0.5%), low-frequency allele (0.5% ≤ AF ≤ 5%), and common allele (AF > 5%) are from left to right. The x-axis represents the functional categories, while the y-axis indicates the variants count. (A) Missense and Synonymous variants. (B) frameshift insertion, frameshift deletion, non-frameshift insertion, and non-frameshift deletion. (C) Stoploss vs. Stopgain. (D) Pathogenic and likely pathogenic variants annotated by ClinVar.

**Figure S8.**
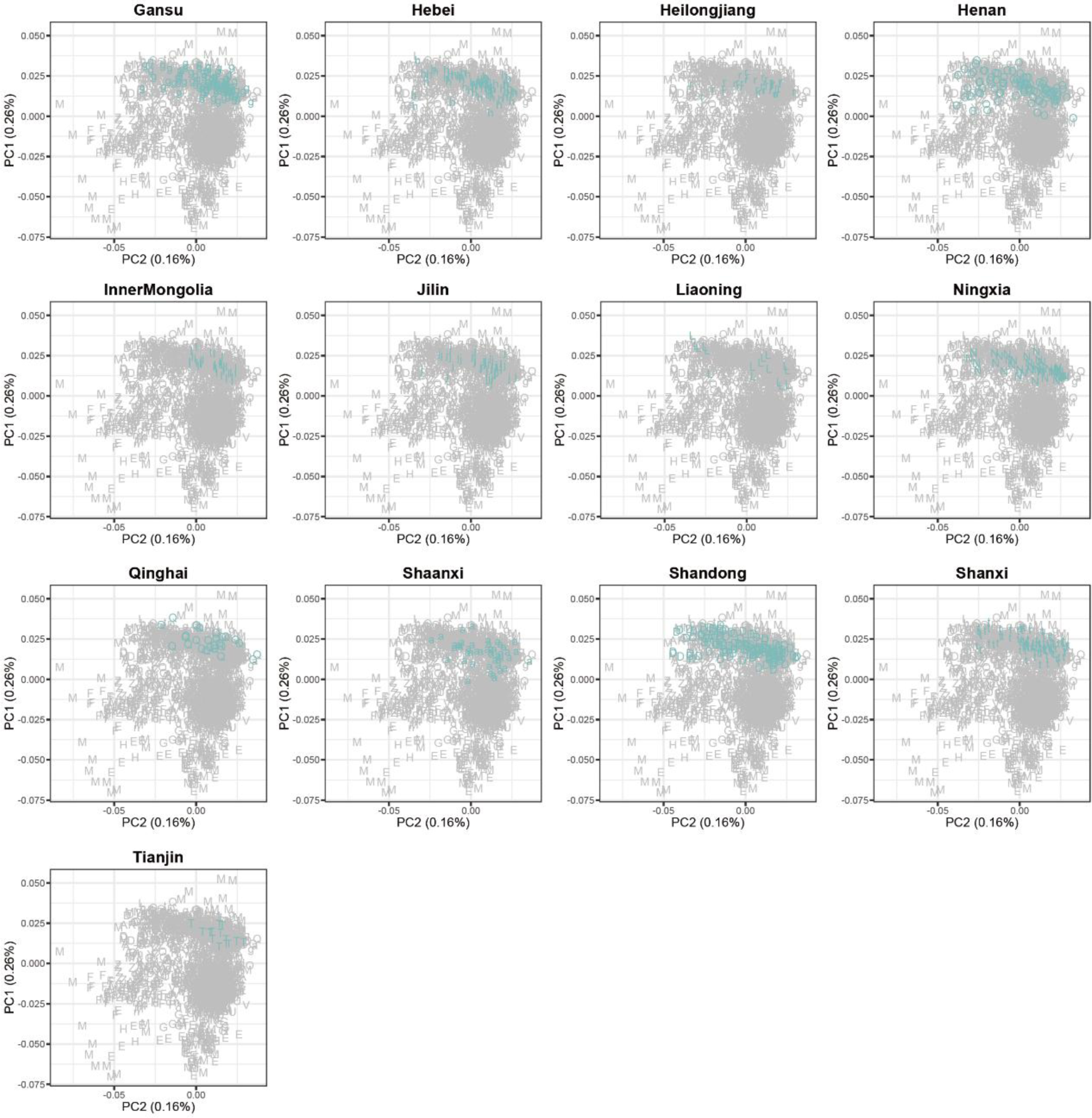
PCA of North Han Chinese samples. Each letter represents one administrative division. The individuals are highlighted in cyan color in north regions, including Gansu (g), Hebei (h), Heilongjiang (r), Henan (O), Inner Mongolia (I), Jilin (j), Liaoning (L), Ningxia (N), Qinghai (Q), Shaanxi (a), Shandong (D), Shanxi (t) and Tianjin (T).

**Figure S9.**
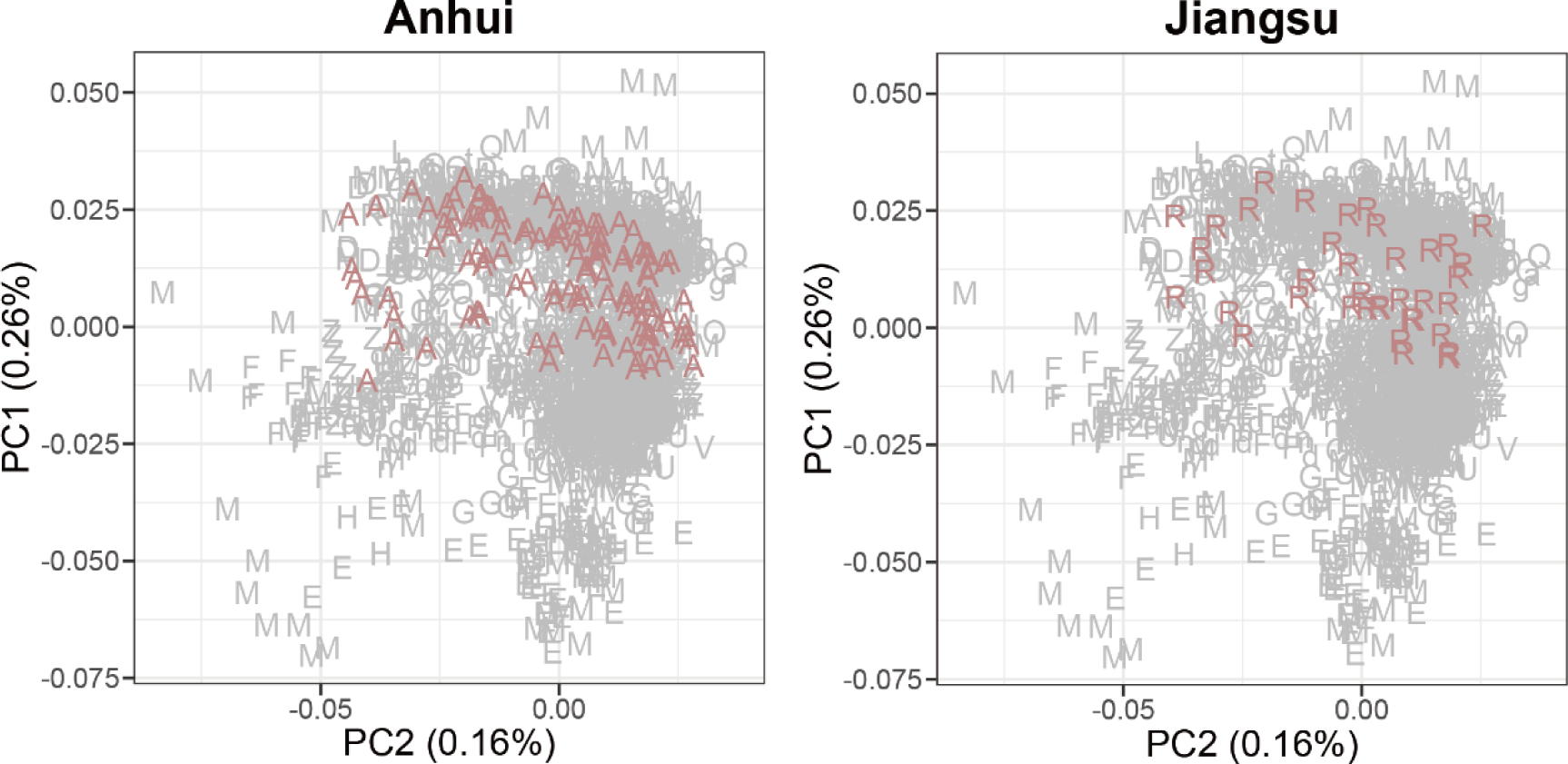
PCA of Central Han Chinese samples. Each letter represents one administrative division. The individuals are highlighted in dark-red for the central regions of Anhui (A) and Jiangsu (R).

**Figure S10.**
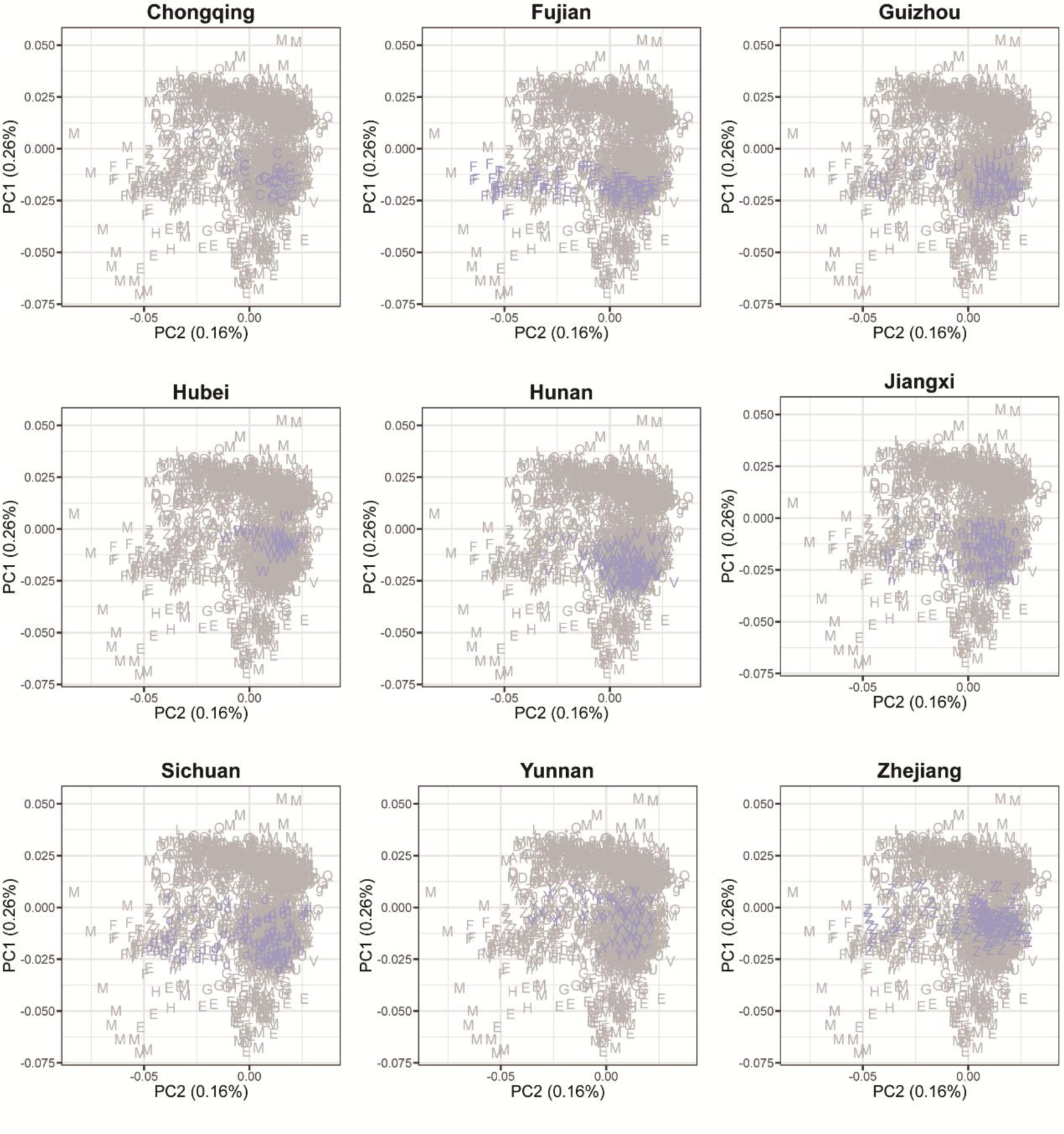
PCA of South Han Chinese samples. Each letter represents one administrative division. The individuals are highlighted in purple c for the southern regions of Chongqing (C), Fujian (F), Guizhou (U), Hubei (W), Hunan (V), Jiangxi (n), Sichuan (d), Yunnan (Y) and Zhejiang (Z).

**Figure S11.**
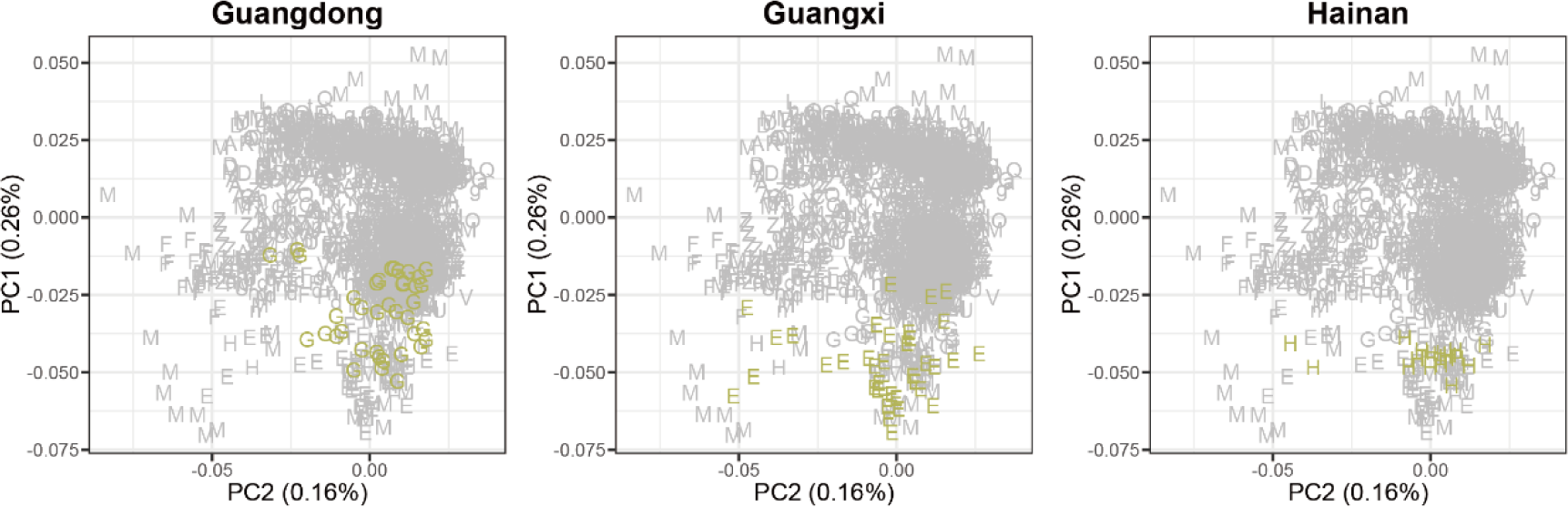
PCA of Lingnan Han Chinese samples. Each letter represents one administrative division. The individuals are highlighted in golden color for the Lingnan regions of Guangxi (E), Guangzhou (G) and Hainan (H).

**Figure S12.**
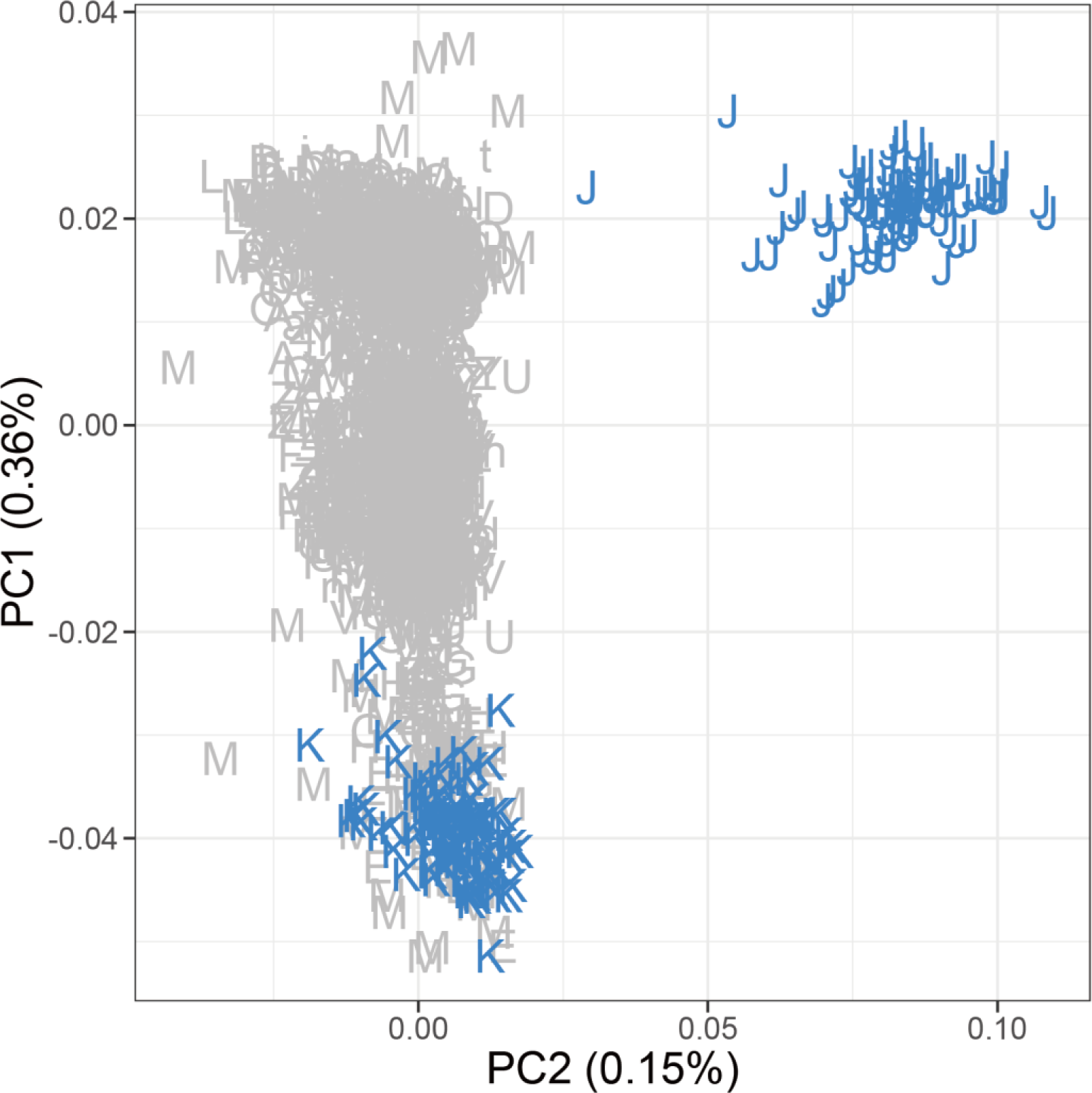
PCA of Chinese, JPT, and KHV individuals from the 1000 Genomes Projects. The blue “J” represents the JPT individuals, while “K” indicates the KHV individuals.

**Figure S13.**
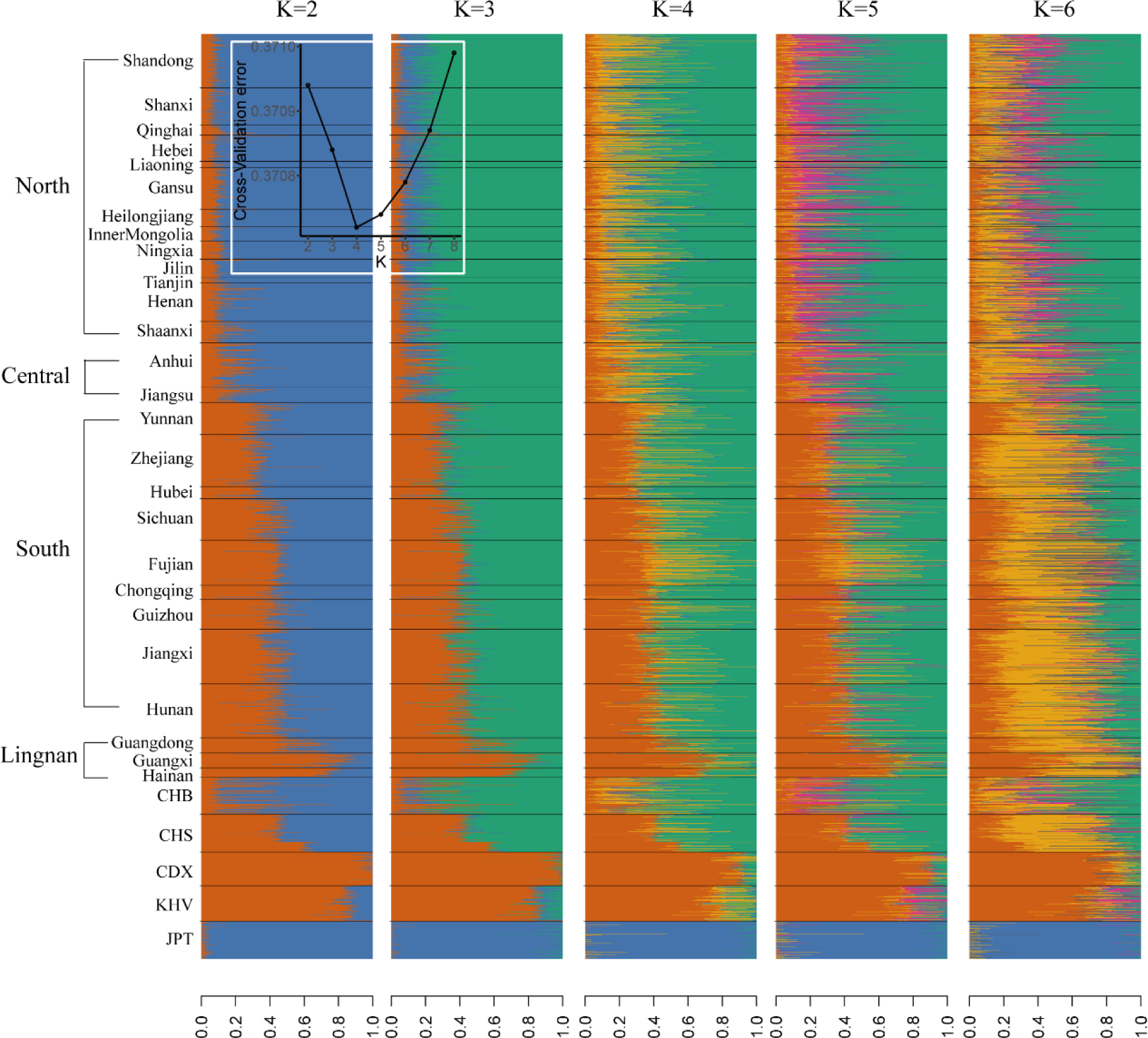
ADMIXTURE analysis of the Han Chinese, CHB, CHS, CDX, KHV and JPT individuals from the 1000 Genomes Projects. The Han Chinese (2,056 individuals) was selected from our WBBC cohort. CHB (103 individuals) is a Han Chinese in Beijing. CHS (105 individuals) is Han Chinese from southern China. CDX (93 individuals) is Chinese Dai in Xishuangbanna. KHV (99 individuals) is Vietnamese from Kinh in Ho Chi Minh City. JPT (104 individuals) is Japanese in Tokyo. The provinces and regions are shown on the left.

**Figure S14.**
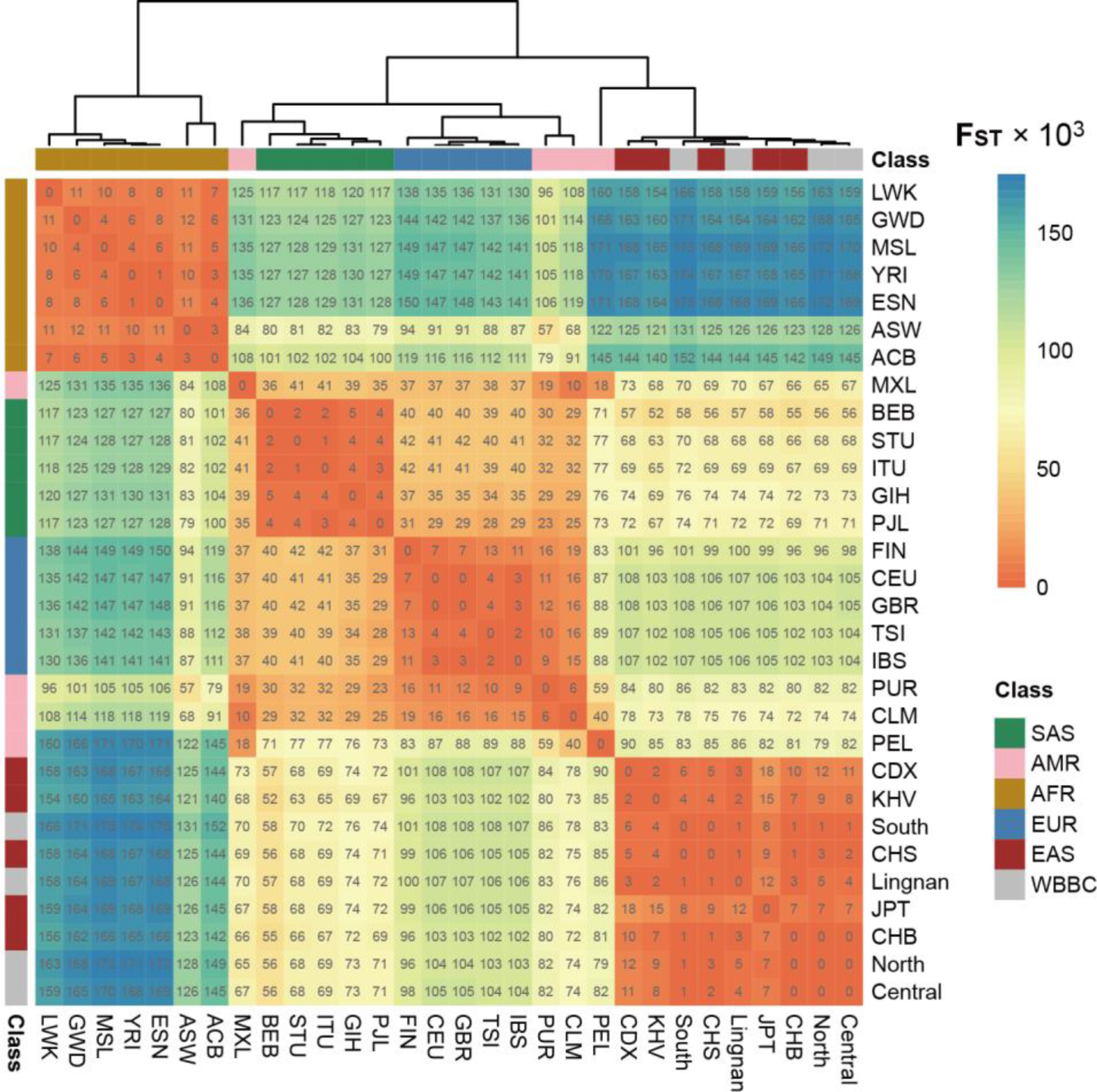
Pairwise F_ST_ between the WBBC and 1KG Project populations. Numbers in each rectangle mean the pairwise F_ST_ value times 1,000. The bars on the top and left show the population classifications. SAS, AMR, AFR, EUR, and EAS are five continent-level ancestry groups of the 1KG Project.

**Figure S15.**
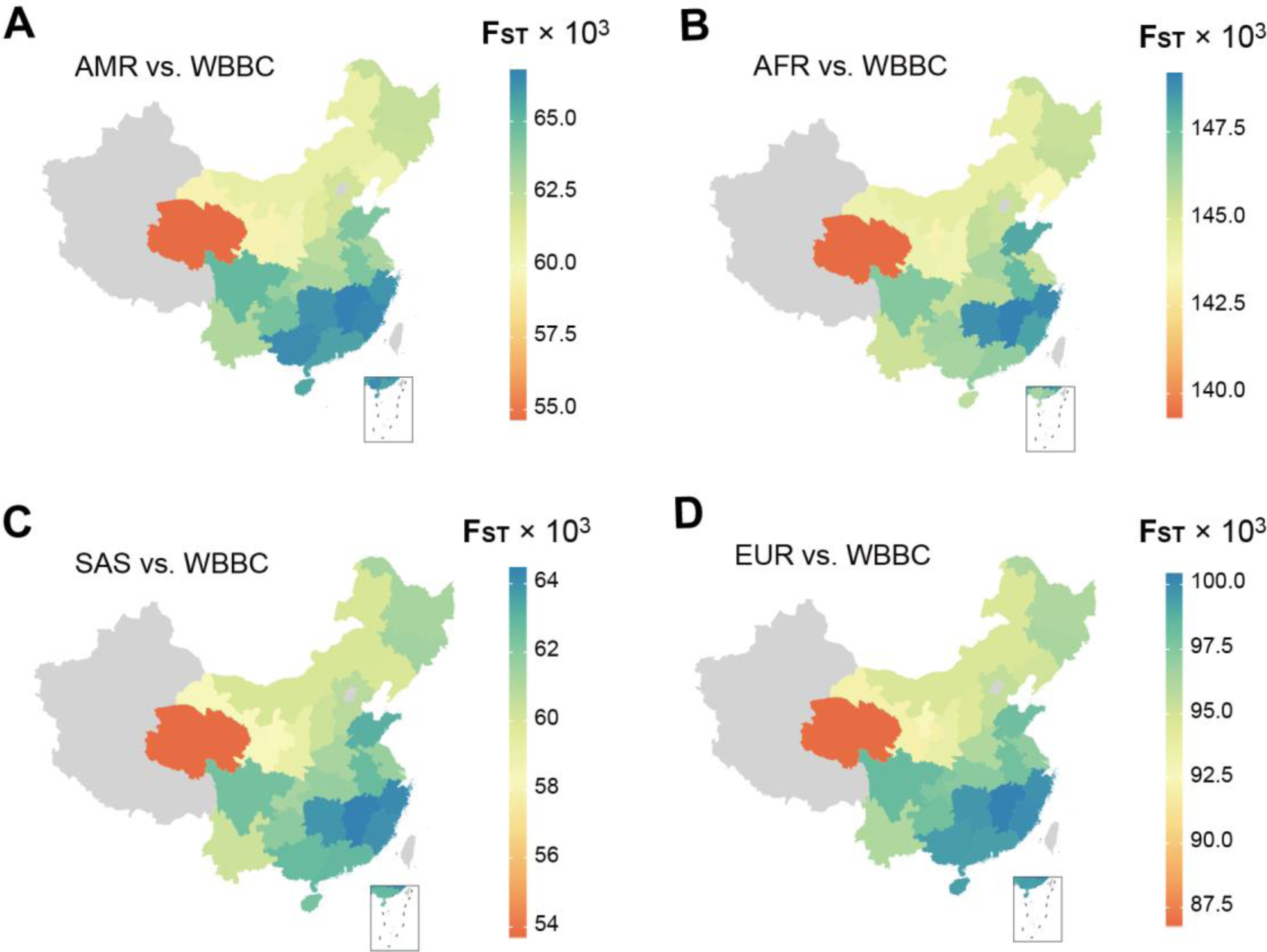
Geographic patterns of pairwise F_ST_ using the 1KG as a reference. Using the four non-Chinese continent-level ancestry groups of the 1KG Project as the reference, including (A) AMR, (B) AFR, (C) SAS, and (D) EUR, we further investigated the geographic patterns of F_ST_ in the 27 administrative divisions. Regions in grey were not sampled.

**Figure S16.**
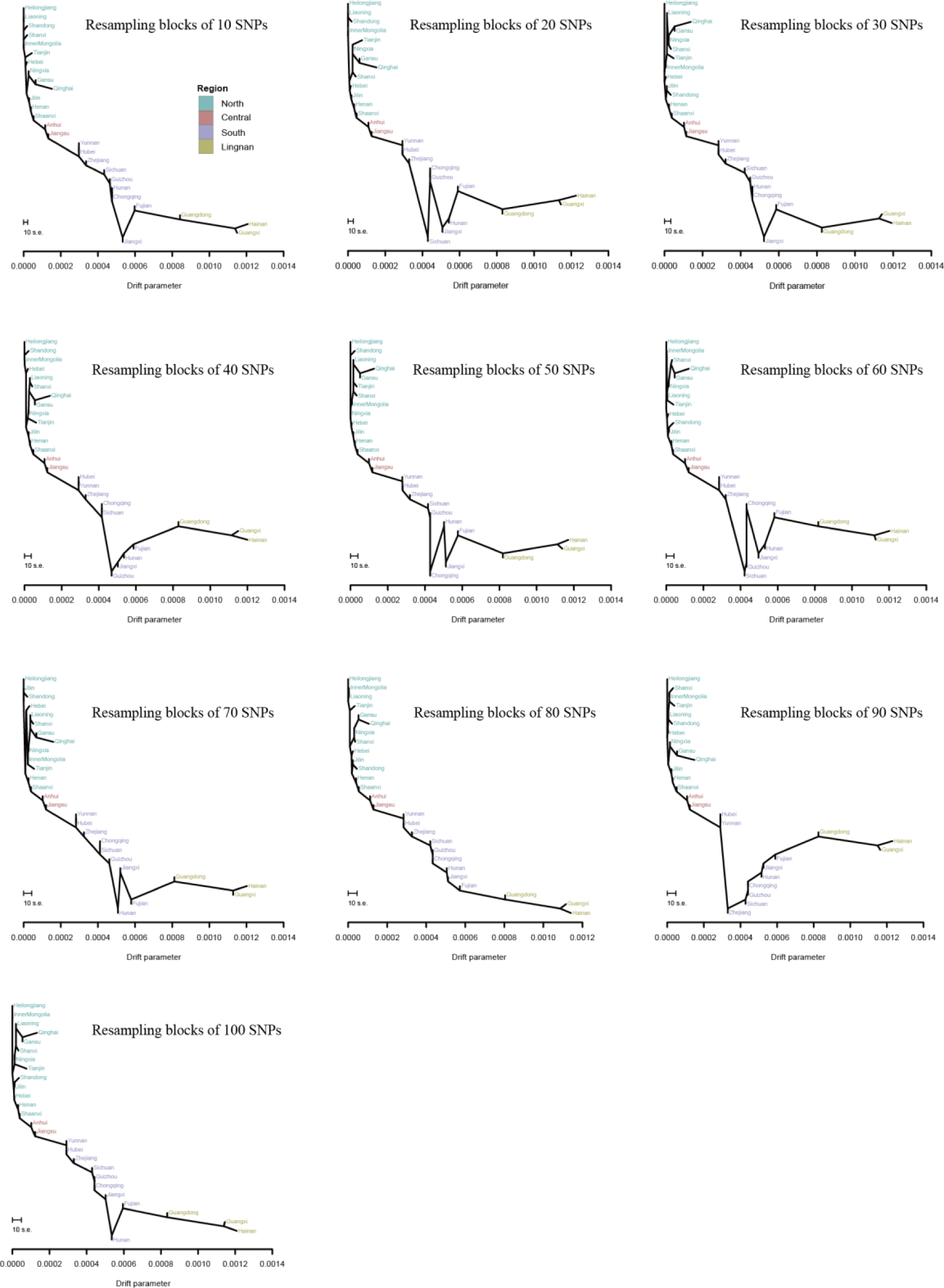
Bootstrap replicates for the tree topology in 27 administrative divisions. The bootstrap replicates were generated by the -bootstrap -k flag of TreeMix software. The plots were y-axis free. The scale bar shows ten times the average standard error of the entries in the sample covariance matrix for the estimated drift parameter.

**Figure S17.**
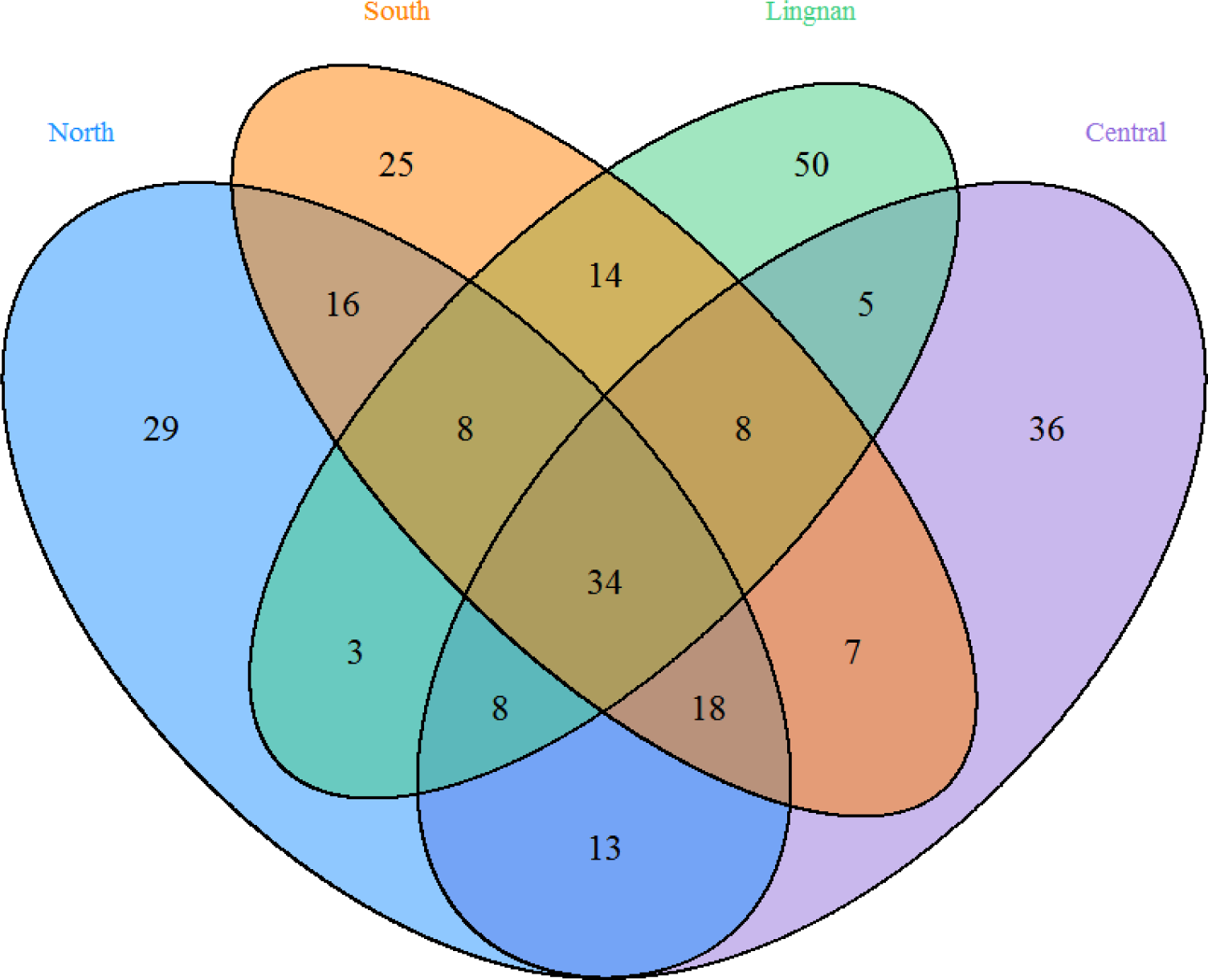
Sharing of genomic regions with higher iHS signals among four subgroups. The number listed within the Venn Diagram indicates the numbers of 200 kb non-overlapping genomic windows in the top 1% of the fraction of SNVs with |iHS| >2.

**Figure S18.**
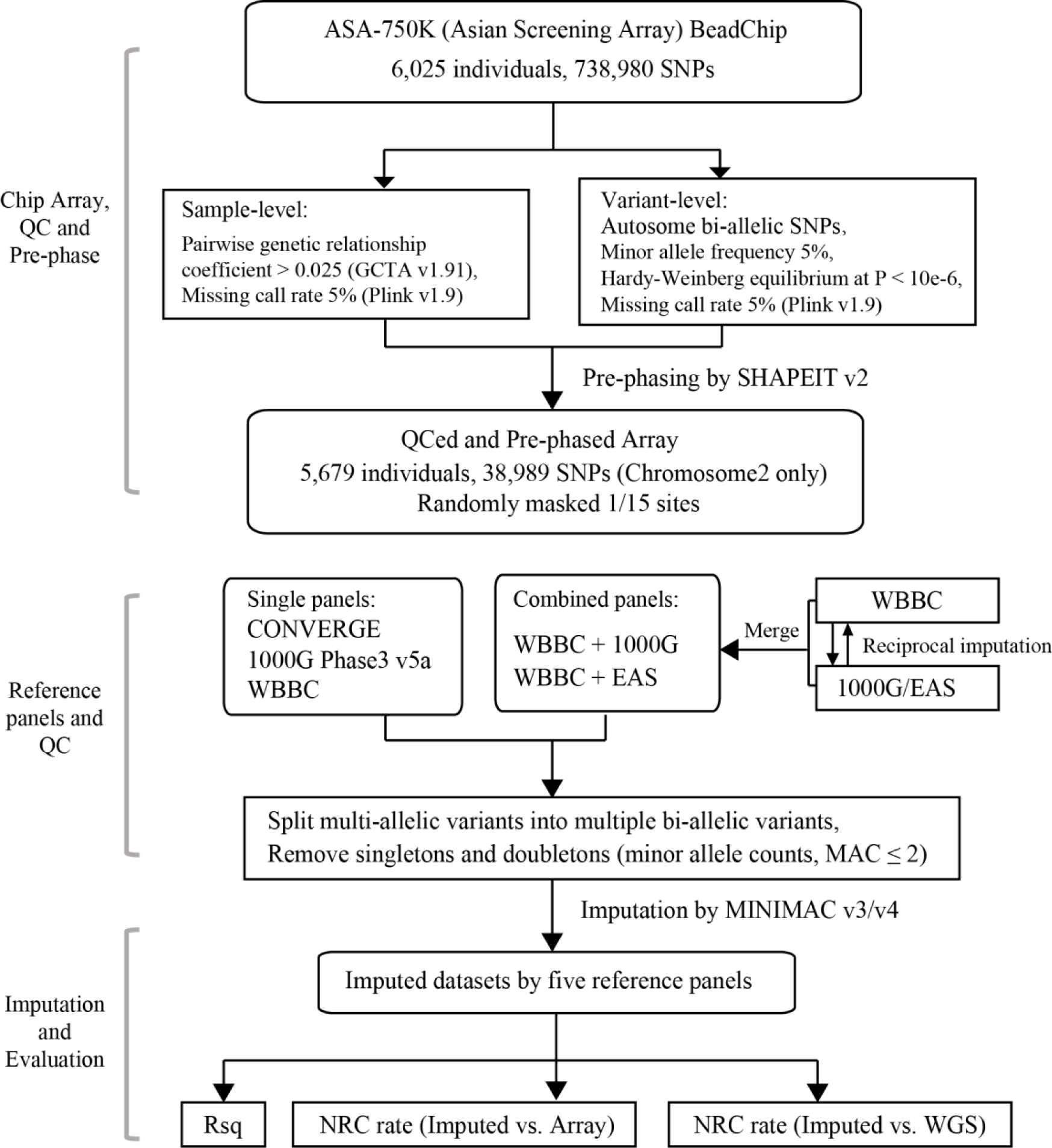
Imputation evaluation workflow. Rsq: R-square value calculated by Minimac4. NRC rate: non-reference allele genotype concordance rate. 1000G included 3,284,591 variants and 5,008 haplotypes; CONVERGE included 1,115,342 variants and 23,340 haplotypes. The WBBC included 2,089,508 variants and 8,610 haplotypes. The WBBC+1000G combined panel consisted of 13,618 haplotypes with 4,450,989 variants. The WBBC+EAS combined panel consisted of 9,618 haplotypes with 2,411,382 variants

**Figure S19.**
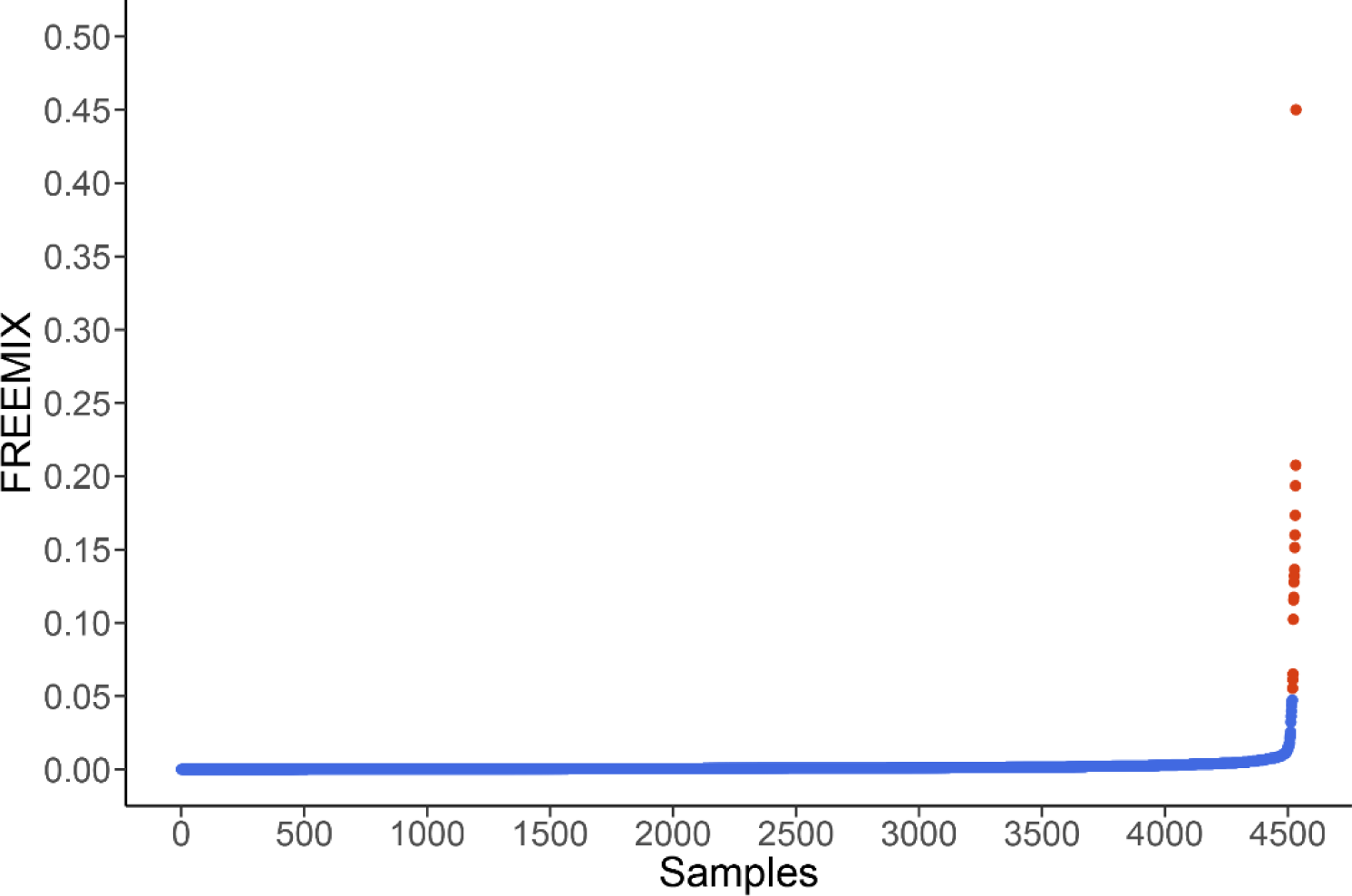
The distribution of the FREEMIX scores was inferred by verifyBamID version1.1.3. The red dots denoted the samples removed by the FREEMIX scores (> 0.05).

**Figure S20.**
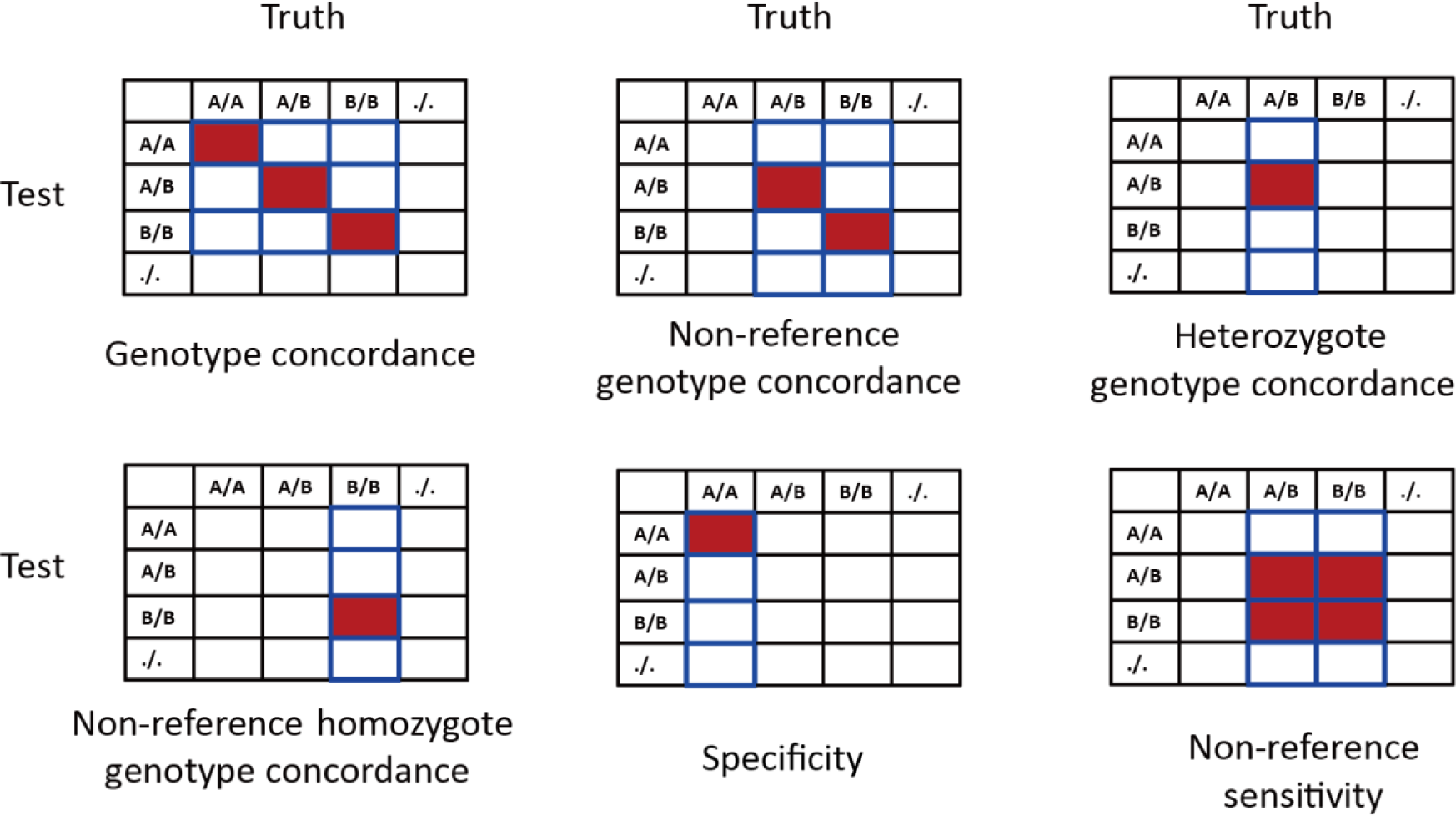
Definition of genotype concordance, specificity, and sensitivity. Letter A represents reference allele while B represents non-reference (NR) allele. The dot means missing called allele. For each metric, the value equals to the corresponding red rectangles divided by all rectangles with a blue border.

**Table S1**. The geographic distribution of all the WBBC cohort samples. The individuals were recruited from 29 of the 34 administrative divisions of the People’s Republic of China (PRC).

**Table S2**. The summary of autosomal variants in each individual in the WBBC cohort.

**Table S3**. The statistics of variants in 4,489 individuals.

**Table S4**. The pathogenic and likely pathogenic variants were recorded by Clinvar in the 1,151 healthy individuals.

**Table S5**. Pair-wise Fst values for 27 provinces of the China in WBBC

**Table S6**. Pair-wise Fst values for 26 populations in the 1KG Phase3 and 4 regions of the China in WBBC

**Table S7**. Pair-wise Fst values for 27 provinces of the China in WBBC and 4 continent groups in the 1KG Phase3

**Table S8**. SNVs with significant selection signatures by SDS analysis in the Han Chinese population.

**Table S9**. The top 1% of non-overlapping genomic windows was identified for positive selection in the North Han Chinese population using the iHS statistic. The adjacent regions, including the same genes, were merged. The clusters were sorted by the fraction of SNVs with |iHS| > 2.

**Table S10**. The top 1% of non-overlapping genomic windows was identified for positive selection in the Central Han Chinese population using the iHS statistic. The adjacent regions, including the same genes, were merged. The clusters were sorted by the fraction of SNVs with |iHS| > 2.

**Table S11**. The top 1% of non-overlapping genomic windows was identified for positive selection in the South Han Chinese population using the iHS statistic. The adjacent regions, including the same genes, were merged. The clusters were sorted by the fraction of SNVs with |iHS| > 2.

**Table S12**. The top 1% of non-overlapping genomic windows was identified for positive selection in the Lingnan Han Chinese population using the iHS statistic. The adjacent regions, including the same genes, were merged. The clusters were sorted by the fraction of SNVs with |iHS| > 2.

**Table S13**. The Kyoto Encyclopedia of Genes and Genomes (KEGG) analysis of candidate genes in the top 1% genomic regions with positive selection by iHS. The terms were selected according to the *p*-value (< 0.05).

## REFERENCES

(2019). Genetics for all. Nat. Genet. 51, 579.

Adzhubei, I., Jordan, D.M., and Sunyaev, S.R. (2013). Predicting functional effect of human missense mutations using PolyPhen-2. Curr Protoc Hum Genet Chapter 7, Unit7 20.

Alexander, D.H., Novembre, J., and Lange, K. (2009). Fast model-based estimation of ancestry in unrelated individuals. Genome Res. 19, 1655–1664.

Bai, W.Y., Zhu, X.W., Cong, P.K., Zhang, X.J., Richards, J.B., and Zheng, H.F. (2019). Genotype imputation and reference panel: a systematic evaluation on haplotype size and diversity. Brief. Bioinform.

Bierut, L.J., Goate, A.M., Breslau, N., Johnson, E.O., Bertelsen, S., Fox, L., Agrawal, A., Bucholz, K.K., Grucza, R., Hesselbrock, V., et al. (2012). ADH1B is associated with alcohol dependence and alcohol consumption in populations of European and African ancestry. Mol. Psychiatry 17, 445–450.

Browning, B.L., and Browning, S.R. (2013). Improving the accuracy and efficiency of identity-by-descent detection in population data. Genetics 194, 459–471.

Browning, B.L., Zhou, Y., and Browning, S.R. (2018). A One-Penny Imputed Genome from Next-Generation Reference Panels. Am. J. Hum. Genet. 103, 338–348.

Cao, Y., Li, L., Xu, M., Feng, Z., Sun, X., Lu, J., Xu, Y., Du, P., Wang, T., Hu, R., et al. (2020). The ChinaMAP analytics of deep whole genome sequences in 10,588 individuals. Cell Res.

Chang, C.C., Chow, C.C., Tellier, L.C., Vattikuti, S., Purcell, S.M., and Lee, J.J. (2015). Second-generation PLINK: rising to the challenge of larger and richer datasets. Gigascience 4, 7.

Chen, J., Zheng, H., Bei, J.X., Sun, L., Jia, W.H., Li, T., Zhang, F., Seielstad, M., Zeng, Y.X., Zhang, X., et al. (2009). Genetic structure of the Han Chinese population revealed by genome-wide SNP variation. Am. J. Hum. Genet. 85, 775–785.

Chiang, C.W.K., Mangul, S., Robles, C., and Sankararaman, S. (2018). A Comprehensive Map of Genetic Variation in the World’s Largest Ethnic Group-Han Chinese. Mol. Biol. Evol. 35, 2736–2750.

Choi, I.G., Son, H.G., Yang, B.H., Kim, S.H., Lee, J.S., Chai, Y.G., Son, B.K., Kee, B.S., Park, B.L., Kim, L.H., et al. (2005). Scanning of genetic effects of alcohol metabolism gene (ADH1B and ADH1C) polymorphisms on the risk of alcoholism. Hum. Mutat. 26, 224–234.

Consortium, U.K., Walter, K., Min, J.L., Huang, J., Crooks, L., Memari, Y., McCarthy, S., Perry, J.R., Xu, C., Futema, M., et al. (2015). The UK10K project identifies rare variants in health and disease. Nature 526, 82–90.

CONVERGE, c. (2015). Sparse whole-genome sequencing identifies two loci for major depressive disorder. Nature 523, 588–591.

Danecek, P., Auton, A., Abecasis, G., Albers, C.A., Banks, E., DePristo, M.A., Handsaker, R.E., Lunter, G., Marth, G.T., Sherry, S.T., et al. (2011). The variant call format and VCFtools. Bioinformatics 27, 2156–2158.

Das, S., Abecasis, G.R., and Browning, B.L. (2018). Genotype Imputation from Large Reference Panels. Annu Rev Genomics Hum Genet 19, 73–96.

Das, S., Forer, L., Schonherr, S., Sidore, C., Locke, A.E., Kwong, A., Vrieze, S.I., Chew, E.Y., Levy, S., McGue, M., et al. (2016). Next-generation genotype imputation service and methods. Nat. Genet. 48, 1284–1287.

Delaneau, O., Marchini, J., and Zagury, J.F. (2011). A linear complexity phasing method for thousands of genomes. Nat Methods 9, 179–181.

Delaneau, O., Zagury, J.F., and Marchini, J. (2013). Improved whole-chromosome phasing for disease and population genetic studies. Nat Methods 10, 5–6.

Druesne-Pecollo, N., Tehard, B., Mallet, Y., Gerber, M., Norat, T., Hercberg, S., and Latino-Martel, P. (2009). Alcohol and genetic polymorphisms: effect on risk of alcohol-related cancer. Lancet Oncol. 10, 173–180.

Edenberg, H.J. (2007). The genetics of alcohol metabolism: role of alcohol dehydrogenase and aldehyde dehydrogenase variants. Alcohol Res Health 30, 5–13.

Ehlers, C.L., Liang, T., and Gizer, I.R. (2012). ADH and ALDH polymorphisms and alcohol dependence in Mexican and Native Americans. Am. J. Drug Alcohol Abuse 38, 389–394.

Eng, M.Y., Luczak, S.E., and Wall, T.L. (2007). ALDH2, ADH1B, and ADH1C genotypes in Asians: a literature review. Alcohol Res Health 30, 22-27.

Eom, S.Y., Hwang, M.S., Lim, J.A., Choi, B.S., Kwon, H.J., Park, J.D., Kim, Y.D., and Kim, H. (2017). Exome-wide association study identifies genetic polymorphisms of C12orf51, MYL2, and ALDH2 associated with blood lead levels in the general Korean population. Environ. Health 16, 11.

Fagerberg, L., Hallstrom, B.M., Oksvold, P., Kampf, C., Djureinovic, D., Odeberg, J., Habuka, M., Tahmasebpoor, S., Danielsson, A., Edlund, K., et al. (2014). Analysis of the human tissue-specific expression by genome-wide integration of transcriptomics and antibody-based proteomics. Mol. Cell. Proteomics 13, 397–406.

Field, Y., Boyle, E.A., Telis, N., Gao, Z., Gaulton, K.J., Golan, D., Yengo, L., Rocheleau, G., Froguel, P., McCarthy, M.I., et al. (2016). Detection of human adaptation during the past 2000 years. Science 354, 760–764.

Fujimoto, A., Kimura, R., Ohashi, J., Omi, K., Yuliwulandari, R., Batubara, L., Mustofa, M.S., Samakkarn, U., Settheetham-Ishida, W., Ishida, T., et al. (2008a). A scan for genetic determinants of human hair morphology: EDAR is associated with Asian hair thickness. Hum. Mol. Genet. 17, 835–843.

Fujimoto, A., Ohashi, J., Nishida, N., Miyagawa, T., Morishita, Y., Tsunoda, T., Kimura, R., and Tokunaga, K. (2008b). A replication study confirmed the EDAR gene to be a major contributor to population differentiation regarding head hair thickness in Asia. Hum. Genet. 124, 179–185.

Gautier, M., Klassmann, A., and Vitalis, R. (2017). rehh 2.0: a reimplementation of the R package rehh to detect positive selection from haplotype structure. Mol. Ecol. Resour. 17, 78–90.

Gelernter, J., Kranzler, H.R., Sherva, R., Almasy, L., Koesterer, R., Smith, A.H., Anton, R., Preuss, U.W., Ridinger, M., Rujescu, D., et al. (2014). Genome-wide association study of alcohol dependence:significant findings in African- and European-Americans including novel risk loci. Mol. Psychiatry 19, 41–49.

Genome of the Netherlands, C. (2014). Whole-genome sequence variation, population structure and demographic history of the Dutch population. Nat. Genet. 46, 818–825.

GenomeAsia, K.C. (2019). The GenomeAsia 100K Project enables genetic discoveries across Asia. Nature 576, 106–111.

Genomes Project, C., Auton, A., Brooks, L.D., Durbin, R.M., Garrison, E.P., Kang, H.M., Korbel, J.O., Marchini, J.L., McCarthy, S., McVean, G.A., et al. (2015). A global reference for human genetic variation. Nature 526, 68–74.

Gudbjartsson, D.F., Helgason, H., Gudjonsson, S.A., Zink, F., Oddson, A., Gylfason, A., Besenbacher, S., Magnusson, G., Halldorsson, B.V., Hjartarson, E., et al. (2015). Large-scale whole-genome sequencing of the Icelandic population. Nat. Genet. 47, 435–444.

Huang, J., Howie, B., McCarthy, S., Memari, Y., Walter, K., Min, J.L., Danecek, P., Malerba, G., Trabetti, E., Zheng, H.F., et al. (2015). Improved imputation of low-frequency and rare variants using the UK10K haplotype reference panel. Nat Commun 6, 8111.

Huang, L., Li, Y., Singleton, A.B., Hardy, J.A., Abecasis, G., Rosenberg, N.A., and Scheet, P. (2009). Genotype-imputation accuracy across worldwide human populations. Am. J. Hum. Genet. 84, 235–250.

Igarashi, M., Nogawa, S., Kawafune, K., Hachiya, T., Takahashi, S., Jia, H., Saito, K., and Kato, H. (2019). Identification of the 12q24 locus associated with fish intake frequency by genome-wide meta-analysis in Japanese populations. Genes Nutr. 14, 21.

Jeon, S., Bhak, Y., Choi, Y., Jeon, Y., Kim, S., Jang, J., Jang, J., Blazyte, A., Kim, C., Kim, Y., et al. (2020). Korean Genome Project: 1094 Korean personal genomes with clinical information. Sci Adv 6, eaaz7835.

Jun, G., Flickinger, M., Hetrick, K.N., Romm, J.M., Doheny, K.F., Abecasis, G.R., Boehnke, M., and Kang, H.M. (2012). Detecting and estimating contamination of human DNA samples in sequencing and array-based genotype data. Am. J. Hum. Genet. 91, 839–848.

Kim, J.W., Choe, Y.M., Shin, J.G., Park, B.L., Shin, H.D., Choi, I.G., and Lee, B.C. (2019). Associations of BRAP polymorphisms with the risk of alcohol dependence and scores on the Alcohol Use Disorders Identification Test. Neuropsychiatr. Dis. Treat. 15, 83–94.

Kowalski, M.H., Qian, H., Hou, Z., Rosen, J.D., Tapia, A.L., Shan, Y., Jain, D., Argos, M., Arnett, D.K., Avery, C., et al. (2019). Use of >100,000 NHLBI Trans-Omics for Precision Medicine (TOPMed) Consortium whole genome sequences improves imputation quality and detection of rare variant associations in admixed African and Hispanic/Latino populations. PLoS Genet. 15, e1008500.

Kumar, P., Henikoff, S., and Ng, P.C. (2009). Predicting the effects of coding non-synonymous variants on protein function using the SIFT algorithm. Nat. Protoc. 4, 1073–1081.

Lander, E.S., and Schork, N.J. (1994). Genetic dissection of complex traits. Science 265, 2037–2048.

Landrum, M.J., Lee, J.M., Riley, G.R., Jang, W., Rubinstein, W.S., Church, D.M., and Maglott, D.R. (2014). ClinVar: public archive of relationships among sequence variation and human phenotype. Nucleic Acids Res. 42, D980–985.

Lek, M., Karczewski, K.J., Minikel, E.V., Samocha, K.E., Banks, E., Fennell, T., O’Donnell-Luria, A.H., Ware, J.S., Hill, A.J., Cummings, B.B., et al. (2016). Analysis of protein-coding genetic variation in 60,706 humans. Nature 536, 285–291.

Levy, D., Ehret, G.B., Rice, K., Verwoert, G.C., Launer, L.J., Dehghan, A., Glazer, N.L., Morrison, A.C., Johnson, A.D., Aspelund, T., et al. (2009). Genome-wide association study of blood pressure and hypertension. Nat. Genet. 41, 677–687.

Li, H. (2011). A statistical framework for SNP calling, mutation discovery, association mapping and population genetical parameter estimation from sequencing data. Bioinformatics 27, 2987–2993.

Li, H., and Durbin, R. (2010). Fast and accurate long-read alignment with Burrows-Wheeler transform. Bioinformatics 26, 589–595.

Li, H., Handsaker, B., Wysoker, A., Fennell, T., Ruan, J., Homer, N., Marth, G., Abecasis, G., Durbin, R., and Genome Project Data Processing, S. (2009). The Sequence Alignment/Map format and SAMtools. Bioinformatics 25, 2078-2079.

Linderman, M.D., Brandt, T., Edelmann, L., Jabado, O., Kasai, Y., Kornreich, R., Mahajan, M., Shah, H., Kasarskis, A., and Schadt, E.E. (2014). Analytical validation of whole exome and whole genome sequencing for clinical applications. BMC Med. Genomics 7, 20.

Liu, S., Huang, S., Chen, F., Zhao, L., Yuan, Y., Francis, S.S., Fang, L., Li, Z., Lin, L., Liu, R., et al. (2018). Genomic Analyses from Non-invasive Prenatal Testing Reveal Genetic Associations, Patterns of Viral Infections, and Chinese Population History. Cell 175, 347–359 e314.

Manichaikul, A., Mychaleckyj, J.C., Rich, S.S., Daly, K., Sale, M., and Chen, W.M. (2010). Robust relationship inference in genome-wide association studies. Bioinformatics 26, 2867–2873.

Martin, A.R., Kanai, M., Kamatani, Y., Okada, Y., Neale, B.M., and Daly, M.J. (2019). Clinical use of current polygenic risk scores may exacerbate health disparities. Nat. Genet. 51, 584–591.

McCarthy, S., Das, S., Kretzschmar, W., Delaneau, O., Wood, A.R., Teumer, A., Kang, H.M., Fuchsberger, C., Danecek, P., Sharp, K., et al. (2016). A reference panel of 64,976 haplotypes for genotype imputation. Nat. Genet. 48, 1279–1283.

McKenna, A., Hanna, M., Banks, E., Sivachenko, A., Cibulskis, K., Kernytsky, A., Garimella, K., Altshuler, D., Gabriel, S., Daly, M., et al. (2010). The Genome Analysis Toolkit: a MapReduce framework for analyzing next-generation DNA sequencing data. Genome Res. 20, 1297–1303.

McVean, G. (2009). A genealogical interpretation of principal components analysis. PLoS Genet. 5, e1000686.

Menozzi, P., Piazza, A., and Cavalli-Sforza, L. (1978). Synthetic maps of human gene frequencies in Europeans. Science 201, 786–792.

Meyer, D., and Thomson, G. (2001). How selection shapes variation of the human major histocompatibility complex: a review. Ann. Hum. Genet. 65, 1–26.

Mou, C., Thomason, H.A., Willan, P.M., Clowes, C., Harris, W.E., Drew, C.F., Dixon, J., Dixon, M.J., and Headon, D.J. (2008). Enhanced ectodysplasin-A receptor (EDAR) signaling alters multiple fiber characteristics to produce the East Asian hair form. Hum. Mutat. 29, 1405–1411.

Nagasaki, M., Yasuda, J., Katsuoka, F., Nariai, N., Kojima, K., Kawai, Y., Yamaguchi-Kabata, Y., Yokozawa, J., Danjoh, I., Saito, S., et al. (2015). Rare variant discovery by deep whole-genome sequencing of 1,070 Japanese individuals. Nat Commun 6, 8018.

Nielsen, R., Akey, J.M., Jakobsson, M., Pritchard, J.K., Tishkoff, S., and Willerslev, E. (2017). Tracing the peopling of the world through genomics. Nature 541, 302–310.

Okada, Y., Momozawa, Y., Sakaue, S., Kanai, M., Ishigaki, K., Akiyama, M., Kishikawa, T., Arai, Y., Sasaki, T., Kosaki, K., et al. (2018). Deep whole-genome sequencing reveals recent selection signatures linked to evolution and disease risk of Japanese. Nat Commun 9, 1631.

Patterson, N., Moorjani, P., Luo, Y., Mallick, S., Rohland, N., Zhan, Y., Genschoreck, T., Webster, T., and Reich, D. (2012). Ancient admixture in human history. Genetics 192, 1065–1093.

Pickrell, J.K., Coop, G., Novembre, J., Kudaravalli, S., Li, J.Z., Absher, D., Srinivasan, B.S., Barsh, G.S., Myers, R.M., Feldman, M.W., et al. (2009). Signals of recent positive selection in a worldwide sample of human populations. Genome Res. 19, 826–837.

Pickrell, J.K., and Pritchard, J.K. (2012). Inference of population splits and mixtures from genome-wide allele frequency data. PLoS Genet. 8, e1002967.

Popejoy, A.B., and Fullerton, S.M. (2016). Genomics is failing on diversity. Nature 538, 161–164.

Price, A.L., Patterson, N.J., Plenge, R.M., Weinblatt, M.E., Shadick, N.A., and Reich, D. (2006). Principal components analysis corrects for stratification in genome-wide association studies. Nat. Genet. 38, 904–909.

Riddell, J., Basu Mallick, C., Jacobs, G.S., Schoenebeck, J.J., and Headon, D.J. (2020). Characterisation of a second gain of function EDAR variant, encoding EDAR380R, in East Asia. Eur. J. Hum. Genet.

Schmidt-Ullrich, R., Aebischer, T., Hulsken, J., Birchmeier, W., Klemm, U., and Scheidereit, C. (2001). Requirement of NF-kappaB/Rel for the development of hair follicles and other epidermal appendices. Development 128, 3843–3853.

Schwarz, J.M., Cooper, D.N., Schuelke, M., and Seelow, D. (2014). MutationTaster2: mutation prediction for the deep-sequencing age. Nat Methods 11, 361–362.

Sherry, S.T., Ward, M.H., Kholodov, M., Baker, J., Phan, L., Smigielski, E.M., and Sirotkin, K. (2001). dbSNP: the NCBI database of genetic variation. Nucleic Acids Res. 29, 308–311.

Shi, Y., Li, L., Wang, Y., Chen, J., and Stanley, H.E. (2019). A study of Chinese regional hierarchical structure based on surnames. Physica A 518, 169–176.

Sirugo, G., Williams, S.M., and Tishkoff, S.A. (2019). The Missing Diversity in Human Genetic Studies. Cell 177, 1080.

Tan, J., Yang, Y., Tang, K., Sabeti, P.C., Jin, L., and Wang, S. (2013). The adaptive variant EDARV370A is associated with straight hair in East Asians. Hum. Genet. 132, 1187–1191.

Terhorst, J., Kamm, J.A., and Song, Y.S. (2017). Robust and scalable inference of population history from hundreds of unphased whole genomes. Nat. Genet. 49, 303–309.

Thayer, T., Arora, A., Batai, K., Halladay, S., Lutz, K., Coleman, A., Pauciulo, M., Kittles, R., Garcia, J.G.N., Yuan, J.X.J., et al. (2020). Sorting Nexin 29 (SNX29) as a Novel Biomarker for Vasoresponsive Pulmonary Arterial Hypertension. Am. J. Respir. Crit. Care Med. 201, A4397–A4397.

Timpson, N.J., Greenwood, C.M.T., Soranzo, N., Lawson, D.J., and Richards, J.B. (2018). Genetic architecture: the shape of the genetic contribution to human traits and disease. Nat Rev Genet 19, 110–124.

Van der Auwera, G.A., Carneiro, M.O., Hartl, C., Poplin, R., Del Angel, G., Levy-Moonshine, A., Jordan, T., Shakir, K., Roazen, D., Thibault, J., et al. (2013). From FastQ data to high confidence variant calls: the Genome Analysis Toolkit best practices pipeline. Curr Protoc Bioinformatics 43, 11 10 11-11 10 33.

Voight, B.F., Kudaravalli, S., Wen, X., and Pritchard, J.K. (2006). A map of recent positive selection in the human genome. PLoS Biol. 4, e72.

Wang, K., Li, M., and Hakonarson, H. (2010). ANNOVAR: functional annotation of genetic variants from high-throughput sequencing data. Nucleic Acids Res. 38, e164.

Weir, B.S., and Cockerham, C.C. (1984). Estimating F-Statistics for the Analysis of Population Structure. Evolution 38, 1358-1370.

Wilcoxin, F. (1947). Probability tables for individual comparisons by ranking methods. Biometrics 3, 119–122.

Wu, D., Dou, J., Chai, X., Bellis, C., Wilm, A., Shih, C.C., Soon, W.W.J., Bertin, N., Lin, C.B., Khor, C.C., et al. (2019). Large-Scale Whole-Genome Sequencing of Three Diverse Asian Populations in Singapore. Cell 179, 736–749 e715.

Xie, G., Lin, Q., Wu, Y., and Hu, Z. (2020). The Late Paleolithic industries of southern China (Lingnan region). Quaternary International 535, 21–28.

Xu, S., Yin, X., Li, S., Jin, W., Lou, H., Yang, L., Gong, X., Wang, H., Shen, Y., Pan, X., et al. (2009). Genomic dissection of population substructure of Han Chinese and its implication in association studies. Am. J. Hum. Genet. 85, 762–774.

Yang, J., Lee, S.H., Goddard, M.E., and Visscher, P.M. (2011). GCTA: a tool for genome-wide complex trait analysis. Am. J. Hum. Genet. 88, 76–82.

Yu, G., Wang, L.G., Han, Y., and He, Q.Y. (2012). clusterProfiler: an R package for comparing biological themes among gene clusters. OMICS 16, 284–287.

Zhu, X., Liu, K., Wang, P., Liu, J., Chen, J., Xu, X., Xu, J., Qiu, M., Sun, Y., Liu, C., et al. (2020). Cohort profile: The Westlake BioBank for Chinese (WBBC) pilot cohort: a prospective study for the late adolescence. medRxiv, 2020.2012.2016.20248291.

